# Spatiotemporal development of growth and death zones in expanding bacterial colonies driven by emergent nutrient dynamics

**DOI:** 10.1101/2023.08.27.554977

**Authors:** Harish Kannan, Paul Sun, Tolga Çağlar, Pantong Yao, Brian R. Taylor, Kinshuk Sahu, Daotong Ge, Matteo Mori, Mya Warren, David Kleinfeld, JiaJia Dong, Bo Li, Terence Hwa

## Abstract

Bacterial colony growth on hard agar is commonplace in microbiology; yet, what occurs inside a growing colony is complex even in the simplest cases. Robust colony expansion kinetics featuring a linear radial growth and a saturating vertical growth indicates a common developmental program which is elucidated here for *Escherichia coli* cells using a combination of modeling and experiments. Radial colony expansion is found to be limited by mechanical factors rather than nutrients as commonly assumed. In contrast, vertical expansion is limited by glucose depletion inside the colony, an effect compounded by reduced growth yield due to anaerobiosis. Carbon starvation in the colony interior results in substantial cell death within 1-2 days, with a distinct death zone that expands with the growing colony. Overall, the development of simple colonies lacking EPS production and differentiation is dictated by an interplay of mechanical constraints and emergent nutrient gradients arising from obligatory metabolic processes.

## INTRODUCTION

The formation of bacterial colonies from individual cells on hard surfaces is one of the simplest modes of bacterial growth in nature and in the laboratory. While there is much interest and extensive research in understanding the formation of biofilms by EPS-producing and quorum-sensing cells (1–11), there is a lot left to be understood already at the quantitative level for simpler colonies formed by cells lacking such capabilities. Colonies of non-EPS-producing *E. coli* cells growing on hard agar exhibit robust expansion characteristics featuring a constant radial expansion rate, after a few hours post-inoculation, that lasts for a few days (12–15), while the colony height increases approximately linearly with time initially between 12-24 hours before slowing down and eventually saturating (13, 14, 16). The constant radial expansion, also observed in colonies of diverse microorganisms (12, 15, 17, 18), has been attributed to be limited by nutrients in the past (12, 15). The saturation of vertical colony expansion, also observed for various microorganisms has been characterized in great experimental detail recently by Ref. (16). Several studies, both computational and experimental, have provided detailed views of the complex metabolic interactions within the colony involving metabolite cross-feeding (13, 19–22); but these studies do not quantitatively connect how the behavior of individual cells relate to overall colony expansion and development.

The kinetics of early colony growth, referred henceforth as the *establishment phase* of colony development, was previously investigated by several of us computationally using an agent-based model of colony growth (23). It was shown that the linear radial growth of the colony in the establishment phase is not limited by nutrient but by mechanical factors controlled by the interplay between surface tension and cell-agar friction. It was also shown that the vertical growth, which was also linear in the establishment phase, is limited by the upward penetration of nutrients from the agar at the bottom of the colony. However, a number of factors limit the application of the model in Ref. (23) to study colony dynamics beyond the establishment phase, including the simplicity of the single nutrient metabolic model used and computational limitations for probing large colonies.

In this work, we extend the agent-based model of Ref. (23) to study the *post-establishment* phase of colony development. Our numerical simulations point to the depletion of the primary nutrient, glucose, inside the colony as the main cause of vertical slowdown, with this depletion accelerated by the deprivation of oxygen in the colony interior even when glucose concentration is high in the agar region below the colony. Substantial cell death driven by anaerobic carbon starvation in the colony interior is predicted to occur within 1-2 days by the simulations and the predicted death zone is validated by direct experimental measurements. Overall, our study highlights the crucial role played by emergent nutrient gradients in controlling colony expansion while also shedding light on nutrient starvation induced cell death within the colony, a surprisingly underexplored aspect of this very common mode of bacterial growth.

## RESULTS

### Initial glucose concentration affects vertical but not radial expansion

To probe the potential role of nutrient limitation, in particular, carbon limitation, on colony expansion, we characterized the morphology of *E. coli* colonies growing on hard agar minimal media plates, focusing on the effect of the initial glucose concentration. A non-motile *E. coli* K12 strain with constitutive GFP expression (EQ59) was used. Each colony originated from a single cell taken from exponentially growing batch culture in the same growth medium, and the colony seeding density was kept low such that the initial inter-colony distance was no smaller than ∼ 1 cm (see Methods).

Measurements of colony dimensions were performed periodically using confocal microscopy. As previously reported, the colony morphology is radially symmetric (**Extended Data Fig. 1, Supplementary Video 1,2**) and can be represented by the *cross-sectional profile,* i.e., a plot of the radial dimension at various vertical distance from the agar surface, as shown in **Fig. 1A, 1B** for two different initial glucose concentrations. Colonies grown on 10 mM glucose plates (**Fig. 1A, Extended Data Fig. 2F**) appear flatter than identically aged colonies grown on 20 mM glucose plates (**Fig. 1B, Extended Data Fig. 2G**) while their radii, defined as the maximum radial dimension at the agar surface, appear similar. Quantitatively, colony radial expansion was linear in time between 15 h and 50 h for colonies grown on both 10 mM glucose plates (∼ 43.2 μm/h) and 20 mM glucose plates (∼ 46.4 μ m/h) (**Fig. 1C, Extended Data Fig. 2C**). Vertical colony expansion, defined by the changes in the vertical dimension at the colony center (referred to as “colony height”) had a fast expansion rate between 15 h and 25 h (∼5.5 μm/h for 10 mM glucose plates and ∼9.5 μm/h for 20 mM glucose plates (**Extended Data Fig. 2D**)). Beyond 25 h, vertical expansion slowed down, and colonies grown on 20 mM glucose plates reach larger heights than those grown on 10 mM glucose plates (**Fig. 1D, Extended Data Fig. 2D**). These observations are also reflected in the overall volume of the colonies (**Fig. 1E, Extended Data Fig. 2E**), estimated from the microscopy images (See Methods).

**Fig. 1.**
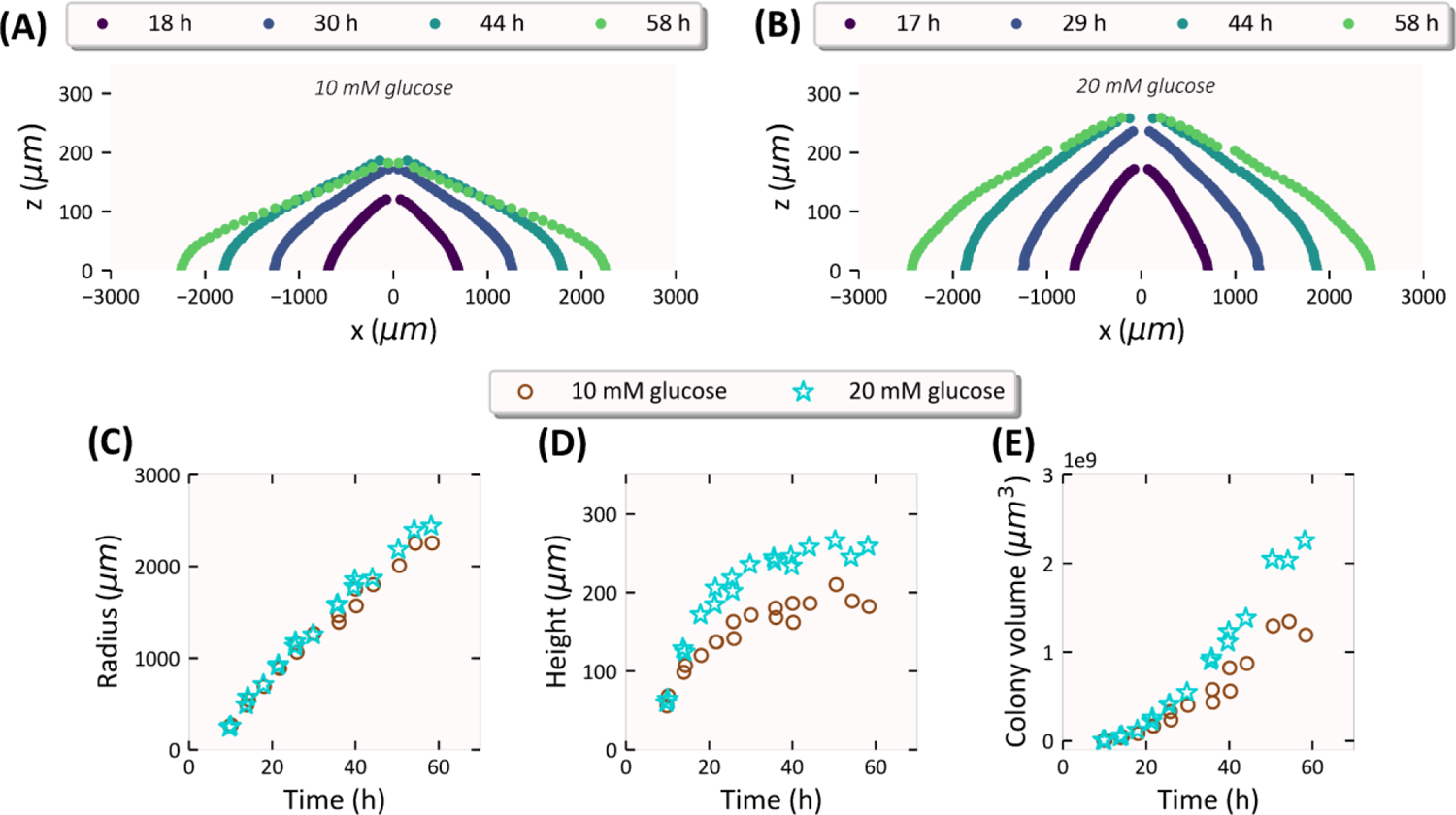
Dependence of E. coli colony expansion on initial glucose concentration. Expansion kinetics of EQ59 *E. coli* colonies on 1.5 %(w/v) agar plates prepared with a defined concentration (10 mM, 20 mM) of glucose, 10 mM ammonium chloride, and 112 mM phosphate buffered saline (PBS) at various times post-inoculation as a single cell. The seeding density of colonies is such that there are ∼10 well-separated colony forming units **(SI. Appendix Fig. S2B)** on a petri dish which is 60 mm diameter and has ∼ 8 mm media depth with a total media volume of ∼ 16 ml. The cross-sectional profile of a colony grown on a minimal media hard agar plate with **(A)** 10 mM glucose and **(B)** 20 mM glucose as carbon source shown for various times (coded by color) post-inoculation. **(C)** The radius (μm), **(D)** The height (μm) and **(E)** The volume (μm^3^) of the colonies plotted against the time (h) post-inoculation. Brown circles represent colonies grown on minimal media plates with 10 mM glucose while cyan stars represent colonies with 20 mM glucose as the carbon source.

### An agent-based reaction-diffusion model captures colony expansion dynamics

To gain insight into the spatiotemporal dynamics of colony expansion, and particularly the role of metabolism as shown in **Fig. 1**, we expanded a previously introduced agent-based model coupled with reaction-diffusion equations to study metabolic interactions due to gradients in the concentrations of key metabolites. Based on several prior studies on microbial biofilms and colony formation (12, 15, 19, 24–29), oxygen is expected to be quickly depleted in the highly dense colony interior. Thus, it is crucial to include both the aerobic and anaerobic growth on glucose in the model for cell metabolism within colony (**Fig. 2A, 2B**). To reduce computational time, we adopted a minimal metabolic model that includes only acetate in addition to glucose and oxygen. Here, acetate represents the bulk of fermentation waste products excreted by *E. coli* during anaerobic growth (**Fig. 2B**). Importantly, acetate can only be metabolized in the presence of oxygen (**Fig. 2C**). Additionally, we included the effect of carbon uptake on cell maintenance (30), the deficit of which, i.e., starvation, leads to cell death (31). Details of the metabolic model are described in **SI Appendix Section 2**. Most of the parameters used are based on experimental measurements as listed in **SI Appendix Table 1**. This metabolic model is coupled to other components of the full model as follows: 1) A discrete, agent-based model of the growth, division, and movement of individual bacterial cells, taking growth rate as the output of the metabolic model (**Fig. 2F**); 2) A system of reaction-diffusion partial differential equations (PDEs) capturing the dynamics of the three metabolites in both colony and agar, driven by diffusion, and local consumption/excretion in various regions of the colony as generated by the metabolic model (**Fig. 2D**). The reaction-diffusion equations are supplemented with appropriate initial and boundary conditions for the agar and colony region, the latter dynamically defined by the location of the cells (**Fig. 2E**). Details of the full model are described in the Methods section and also in **SI Appendix Sections 1,2**.

**Fig. 2.**
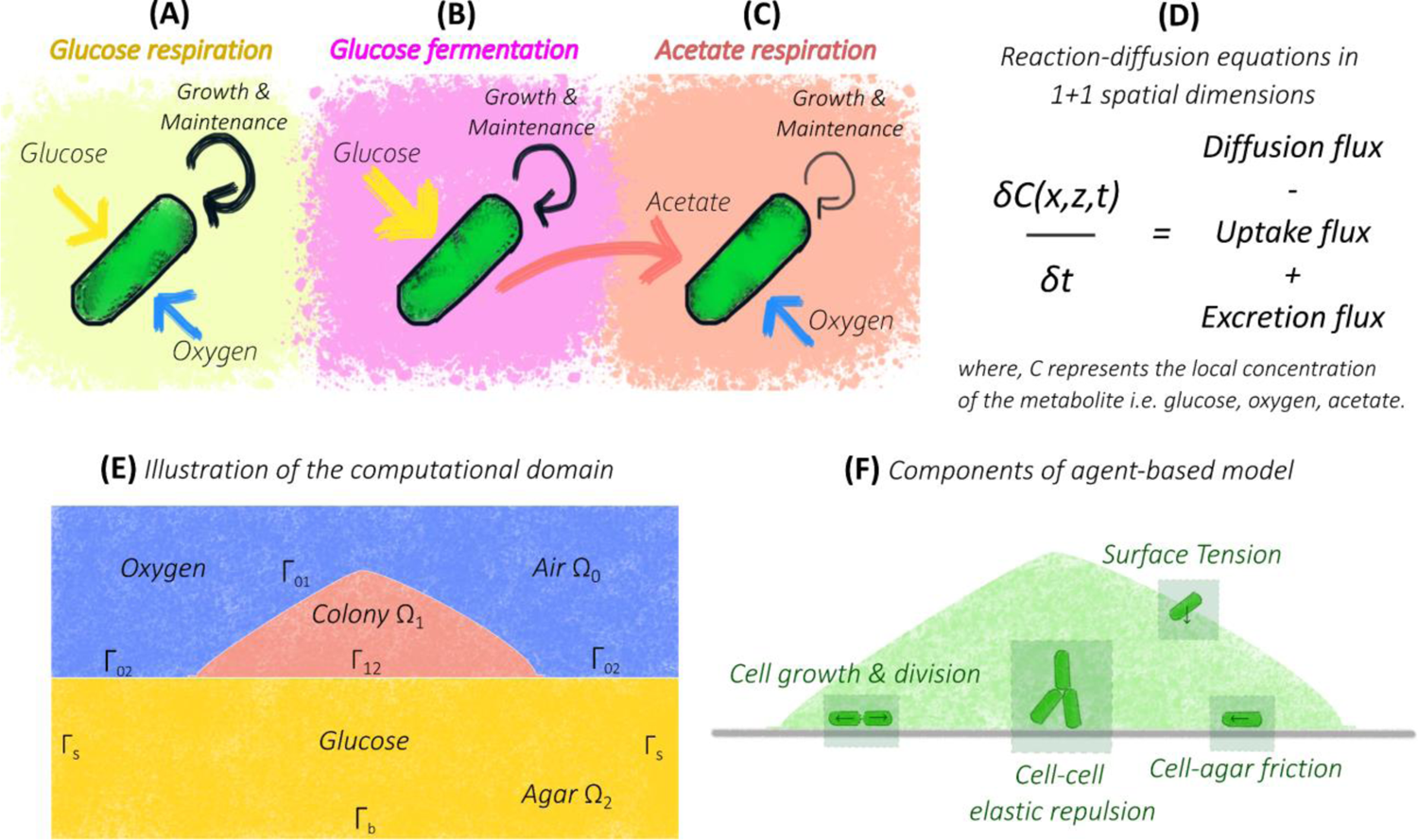
Illustration of the metabolic and agent-based components of model for colony expansion. The different cell growth and maintenance modes included in our model: **(A)** aerobic growth and maintenance on glucose, **(B)** anaerobic growth and maintenance on glucose and **(C)** aerobic growth and maintenance on acetate. Anaerobic consumption of glucose by cells is accompanied by acetate excretion indicated by the red arrow in **(B).** Even under aerobic growth on glucose, cells still excrete a small amount of acetate (not indicated) due to overflow metabolism which is also included in our model. **(D)** A template reaction-diffusion equation used to model the spatiotemporal dynamics of metabolite concentrations. **(E)** Illustration of the computational domain comprising of the colony, agar, and air sub-domains with appropriate boundaries between the sub-domains. **(F)** Illustration of agent-based model features involving individual cell growth, division, and movement. The forces experienced by the cells arise from cell-cell, cell-agar interaction as well as surface tension.

An agent-based model in a three-dimensional setting becomes computationally expensive due to the sheer number of individual cells (roughly 10^9^ cells) that make up a macroscopic bacterial colony. Thus, in this study we instead employ a (1+1)-dimensional model, i.e., the cells occupy a two-dimensional region with one dimension (x) being along the colony-agar interface and the other dimension (z) being the one perpendicular to the agar surface, and with cells allowed to move only in the x-z plane (**SI Appendix Section S3A**). This (1+1)-dimensional setting allows the study of both radial and vertical colony expansion while still enabling the simulations to be computationally tractable.

We performed (1+1)-dimensional simulations of colony growth on top of a rectangular agar region (**Fig. 2E**) with uniform initial glucose and oxygen concentrations, and no acetate. The width and depth of the agar region are chosen to be comparable to the agar cross-section “available” to a colony in our experiments (**SI Appendix Table 2**). The simulations start with a single cell placed at the center of agar-air interface. The cells elongate and replicate due to growth, pushing each other outward and defining a colony region which would expand radially and vertically. The resulting cross-sectional profiles of such expanding colonies are shown in **Fig. 3A, 3B** for two different initial glucose concentrations. Similar to experimental observations (**Fig. 1A, 1B**), the simulated colonies with 10 mM initial glucose concentration (**Fig. 3A**) appear flatter than the colonies with 20 mM glucose (**Fig. 3B**). The increase of colony width (“radius”) is similar for the two cases throughout the duration of the study (∼ 50 h); see **Fig. 3C**. However, the colony height noticeably increases with the initial glucose concentration (**Fig. 3D**): compared to the case with 10 mM initial glucose, the colony grown with 20 mM initial glucose reaches larger heights beyond the first 10 h, along with more cells in the colony (**Fig. 3E**).

**Fig. 3.**
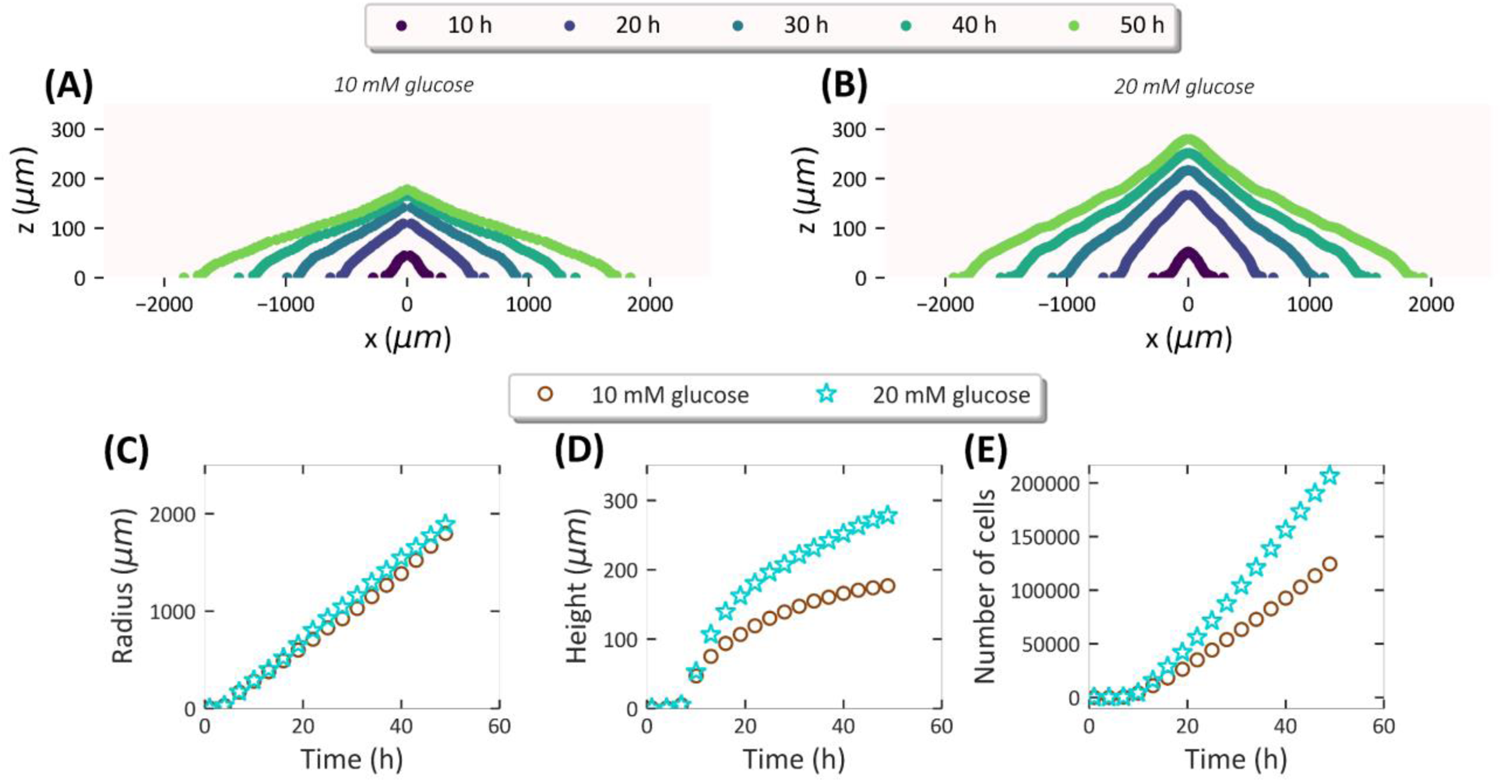
Simulations based on model capture overall colony expansion kinetics. Temporal dynamics of (1+1)-dimensional agent-based simulations of colony expansion for 10 mM initial glucose concentration versus 20 mM initial glucose concentration in agar. Simulations start with a single cell on top of an agar region with dimensions of ∼ 10 mm x 8 mm. The cross-sectional profile of a simulated colony with **(A)** 10 mM initial concentration of glucose in agar and **(B)** 20 mM initial concentration of glucose in agar at various times (coded by color) of colony development. **(C)** The radius (μm), **(D)** The height (μm) and **(E)** The number of cells in the simulated colonies plotted against the time (h) post-inoculation. Brown circles represent simulations with 10 mM initial glucose concentration in agar while cyan stars represent simulations with 20 mM initial glucose concentration in agar.

### Radial colony expansion is not limited by nutrients

Using a single-nutrient (oxygen unlimited) model, Warren et al. (23) showed that during the establishment phase of colony growth it is the orientation of cells that determines the radial expansion rate. In particular, there is the horizontally oriented monolayer of growing cells located at the colony periphery, the width of which determines the magnitude of radial expansion rate. In our model which includes anaerobiosis, the same feature is seen preserved wherein a non-zero horizontal component of velocity is confined to the peripheral monolayer at 15 h and 50 h (**Fig. 4A**). This is a consequence of cells at the periphery being horizontally oriented while becoming progressively vertically oriented towards the center of colony as shown by the snapshot at 15 h (**Extended Data Fig. 4A**). This configuration of cell orientation remains invariant throughout the duration of simulation after 10 h (**Extended Data Fig. 4CD**). Further, the direction of cell movement is highly correlated with cell orientation, with the cells in the peripheral region pointing horizontally outward as shown by a snapshot at 15 h (**Extended Data Fig. 4AE**).

**Fig. 4.**
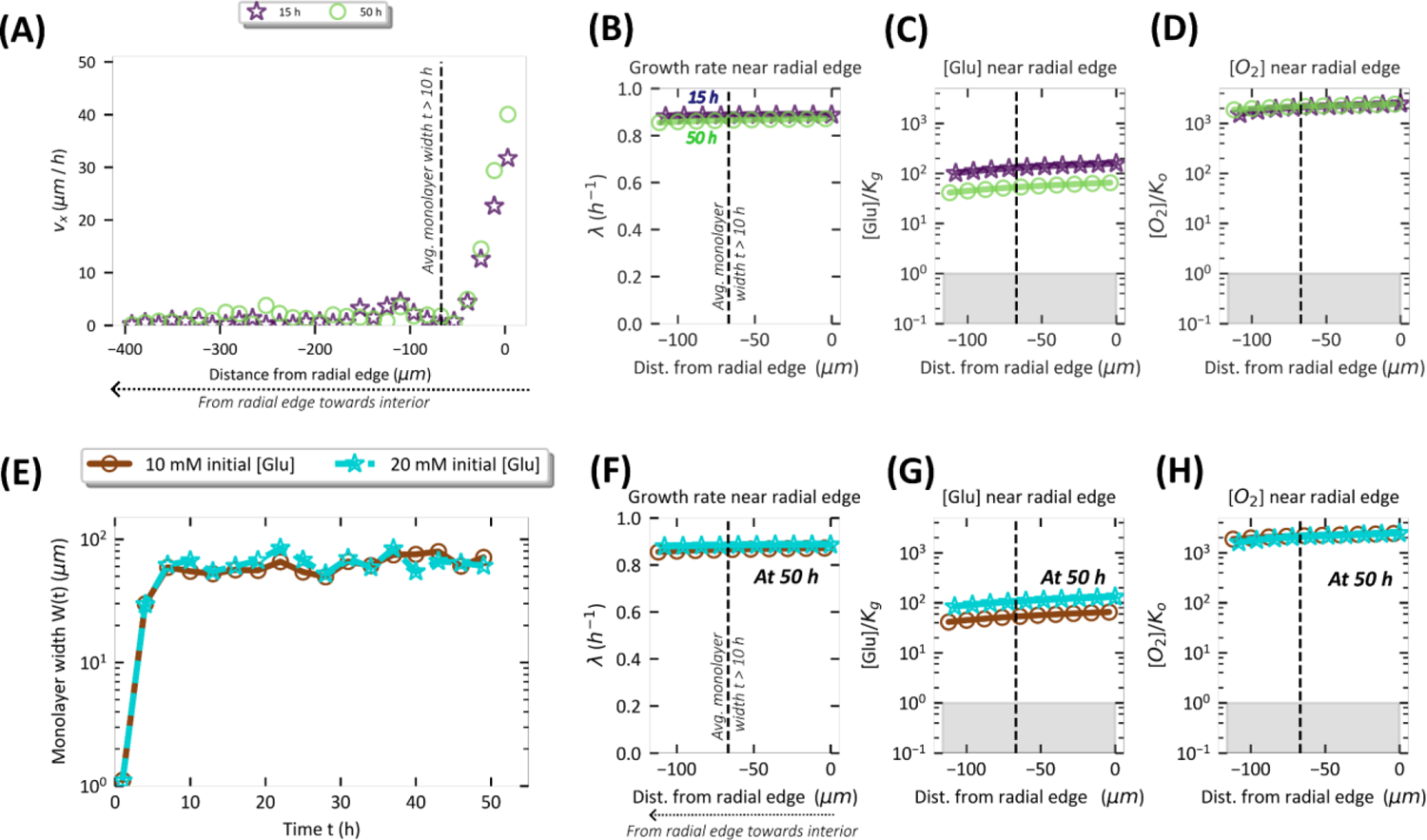
Radial expansion of colony is not limited by nutrients. **(A)** The horizontal velocity component (μm/h) of cells located at the colony-agar interface in simulated colonies plotted as a function of horizontal distance from the peripheral edge of the colony at 15 h and 50 h. A distance of zero represents the radial edge of the colony. The **(B)** growth rate, **(C)** glucose concentration and **(D)** oxygen concentration at the colony-agar interface is plotted against the distance from the radial edge of the colony at 15 h and 50 h of simulated colony development. The nutrient concentrations in **(C)** and **(D)** are normalized by the respective Monod constants and the grey shaded region represents concentration values less than the respective Monod constant. **(A-D)** are for simulations with 10 mM initial glucose concentration and the black dashed vertical line represents the monolayer width averaged for t > 10 h. **(E)** The monolayer width (μm) of the simulated colonies plotted against time for 10 mM initial concentration in agar (brown) and 20 mM initial glucose concentration (cyan) in agar. The **(F)** growth rate, concentration of **(G)** glucose and **(H)** oxygen at the colony-agar interface normalized by the respective Monod constant plotted against the distance from the radial edge of the colony after 50 h of simulated colony development. A distance of zero represents the radial edge of the colony. The grey shaded region represents concentrations below the Monod constant. In **(E-H),** brown symbols represent simulations with 10 mM initial glucose concentration in agar while cyan stars represent simulations with 20 mM initial glucose concentration in agar.

Next, we examine the extent of cell growth which drives cell movement. Cells are seen to grow at close to the maximal rate for at least ∼ 120 μm into the periphery (**Fig. 4B**), well beyond the ∼70 μm monolayer region which determines the radial expansion. This is seen at both 15 h and 50 h of colony development. The near maximal growth rate arises because throughout this region, the glucose and oxygen concentration (**Fig. 4CD**) remain well above the K_g_ and K_o_ value (**SI Appendix Table 1**) respectively during this time period. Thus, our results indicate that radial expansion is limited by neither glucose nor oxygen. Rather, it is limited by the monolayer width which is set by mechanical factors like cell-agar friction and surface tension (23) and not by nutrients. As further supporting evidence, the monolayer width remains similar when the initial glucose concentration is increased from 10 mM to 20 mM (**Fig. 4D**). This explains the independence of radial expansion dynamics on initial glucose concentration during the first 50 h (**Fig. 3C**) since the growth of cells in the monolayer is not limited by nutrients (**Fig. 4FGH**) for either initial condition even though the interior regions are nutrient limited (**Extended Data Fig. 5**).

### Vertical colony expansion is limited by glucose

In contrast to horizontal velocity of cells within colony, the vertical component of cell velocity peaks at the colony center during early stages of development (purple symbols **Fig. 5A**) consistent with the cell orientation profile (**Extended Data Fig. 4**). Over time, the vertical velocity decreases in the colony interior (teal and green symbols **Fig. 5A**), mirroring the slowdown in vertical colony expansion seen in **Fig. 3D**.

**Fig. 5.**
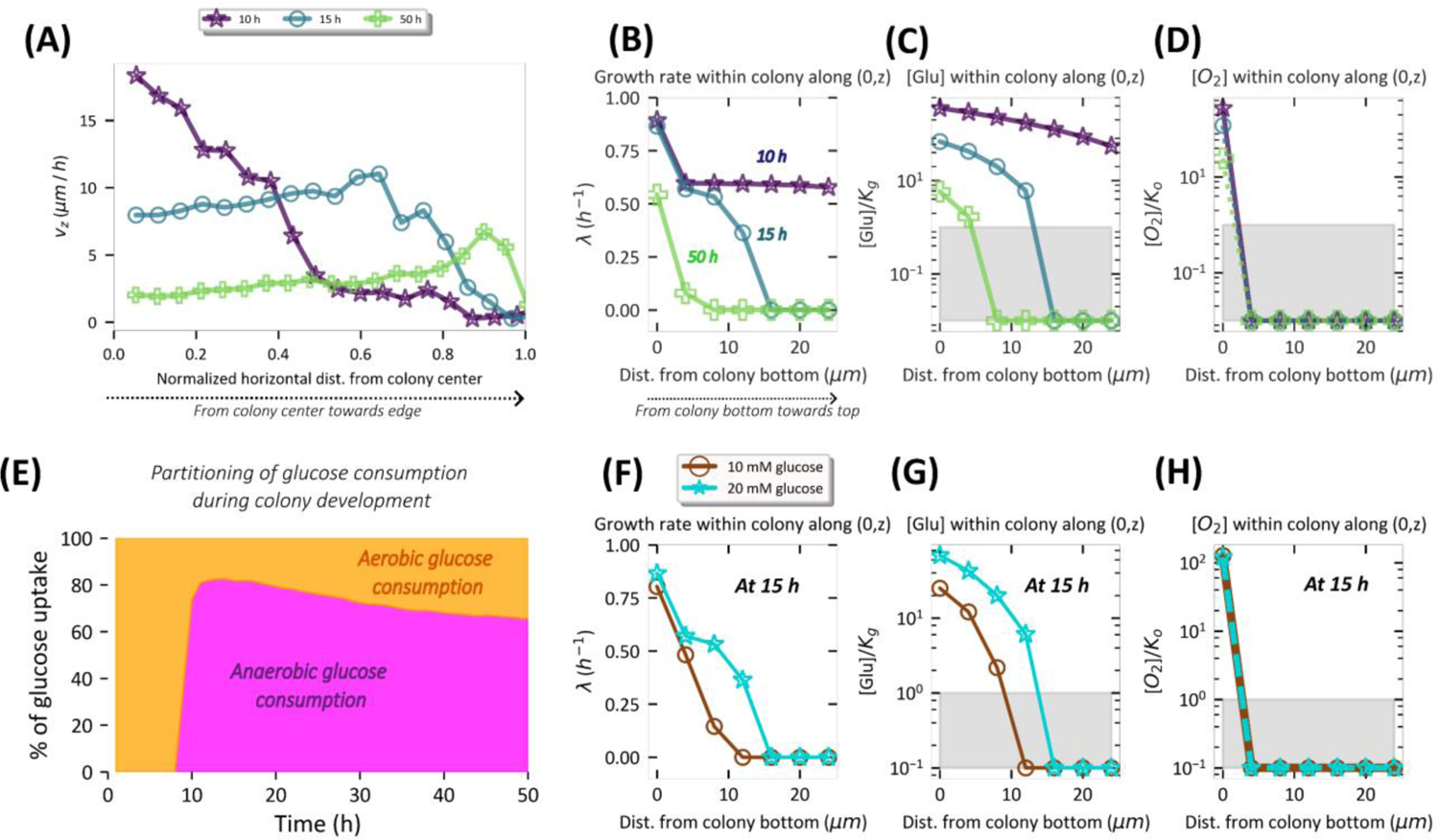
Vertical expansion of colony is limited by glucose. **(A)** The vertical velocity component (μm/h) averaged over the z-coordinate in simulated colonies plotted as a function of horizontal distance from the colony center at 10 h, 15 h and 50 h. A distance of zero represents the colony center. The **(B)** growth rate, **(C)** glucose concentration and **(D)** oxygen concentration along (0,z) axis of colony is plotted against the distance (z) from colony bottom at 10 h, 15 h and 50 h of simulated colony development. The nutrient concentrations in **(C)** and **(D)** are normalized by the respective Monod constants and the grey shaded region represents concentration values less than the respective Monod constant. **(A-D)** are for simulations with 20 mM initial glucose concentration. **(E)** The percentage of glucose uptake flux (integrated over entire colony) towards aerobic (yellow) and anaerobic (magenta) metabolism plotted as a function of time for simulation with 20 mM initial glucose concentration. The **(F)** growth rate, concentration of **(G)** glucose and **(H)** oxygen plotted along (0,z) axis of colony is plotted against the distance (z) from colony bottom at 15 h of simulated colony development. The grey shaded region represents concentrations below the Monod constant. In **(F-H),** brown symbols represent simulations with 10 mM initial glucose concentration in agar while cyan stars represent simulations with 20 mM initial glucose concentration in agar.

To understand the factors underlying the vertical slowdown, we focus on the cells located at the center of the colony (x=0) at various distances from the agar to colony tip (0<z<H). Looking at the growth rate of these cells at the center we find that, while all cells grow at 10 h (**Supplementary Video 6**), the growth at 15 hours is restricted to a narrow regime within ∼20 μm of the agar (**teal symbols Fig. 5B**). Further, by 50 h growth vanishes except for cells at the agar interface. Looking further into the distribution of nutrients required for growth, we see that glucose concentration drops below the Monod constant (**grey region Fig. 5C**) at vertical distances commensurate with the observed drop in growth over time (**Fig. 5B**). On the other hand, oxygen concentration drops below the Monod constant within 4 μm away from agar already at 10 hours (**Fig. 5D**). This indicates that growth after 10 hours in this region is anaerobic. A detailed description of the spatiotemporal dynamics of glucose and oxygen concentration throughout the colony and agar is reported in **Extended Data Fig. 5** and **Supplementary Video 3-4**. It is worth noting that the total amount of glucose in agar is not depleted since glucose consumed is only roughly half of the initial amount even at 50 hours of colony development (**Extended Data Fig. 5A**). Yet, the local concentration within the colony becomes limiting for growth over time; this is a consequence of the high glucose uptake flux due to a combination of the high density of cells (**Extended Data Fig. 4A**) and the high specific uptake rate of glucose during anaerobic growth (**SI appendix Table 1**).

Returning to the dependence of vertical colony expansion on the initial glucose concentration (**Fig. 3D)**, it is seen that 20 mM initial glucose concentration gives rise to a thicker vertical growth zone (∼ 16 μm at 15 h) compared to that for the 10 mM initial glucose concentration (< 10 μm at 15 h) (**Fig. 5F**). A higher initial glucose concentration results in an increased vertical penetration of glucose (**Fig. 5G)** while oxygen remains completely depleted in either case (**Fig. 5H**). This rationalizes the dependence of vertical expansion dynamics on initial glucose concentration reported in **Fig. 3D**.

As shown in **Fig. 5E**, the anaerobic consumption of glucose sharply rises at around 10 hours of colony development. This is a consequence of local oxygen depletion within the colony interior due to consumption of oxygen by cells (**Fig. 5H, Extended Data Fig. 5, *Supplementary Video 4***). Anaerobic consumption of glucose by cells is accompanied by extensive acetate excretion (**Fig. 2B, SI Appendix Table 1**) which can serve as a carbon source for cells elsewhere where oxygen is available, but glucose is not (**Fig. 2C**). (We note that even under aerobic growth on glucose, cells still excrete acetate, albeit at a lower level, due to *overflow metabolism* (32); this feature is also included in our model (**SI Appendix Table 1**).) In our simulations, we see an initial build-up of acetate within the colony to levels of ∼ 8 mM at around 10 h (**Extended Data Fig. 6A**). Acetate concentration drops beyond 10 h, starting from the top of the colony where cells depleted of glucose start to consume acetate (**Extended Data Fig. 6A**). This in turn leads to a vertically decreasing concentration of acetate as the consumption of acetate is highest near the colony-air interface where oxygen is available (**Extended Data Fig. 5E**). The total acetate uptake towards cell growth over time is summarized by the blue points in **Extended Data Fig. 6B**. Also, plotted there is the total acetate uptake towards maintenance (**green stars in Extended Data Fig. 6B**). It is seen that beyond 30 hours, acetate is being predominantly used for cell maintenance rather than for growth (**SI Appendix Videos 7,8**). This arises because when the acetate concentration drops low, maintenance is prioritized over growth (**explained in SI Appendix Section 2**). As a comparison we show in **Extended Data Fig. 6E, Supplementary Video 8**, that consumption of glucose (for both growth and maintenance) is restricted to a few μm above colony-agar interface shortly beyond 10 h.

### The bulk of the colony is under nutrient starvation

Based on our observations of local glucose depletion in the colony bulk, it comes as no surprise that the proportion of growing cells within the simulated colony sharply drops between 10 to 20 hours with the majority of the colony in a non-growing state beyond 20 hours (**Fig. 6A**). This suggests a sub-exponential increase of colony volume beyond ∼ 20h in our experiments (**Fig. 1E, Extended Data Fig. 2E**), which is also captured by the dynamics of the total number of cells within the colony in our simulations (**Fig. 3E**), is a consequence of fewer cells within the colony undergoing growth and division. Thus, it is worth noting that this sub-exponential increase of colony volume is an emergent behavior of our model wherein each cell is modeled such that it only responds to its local environment, specifically, the local concentration of nutrients.

**Fig. 6.**
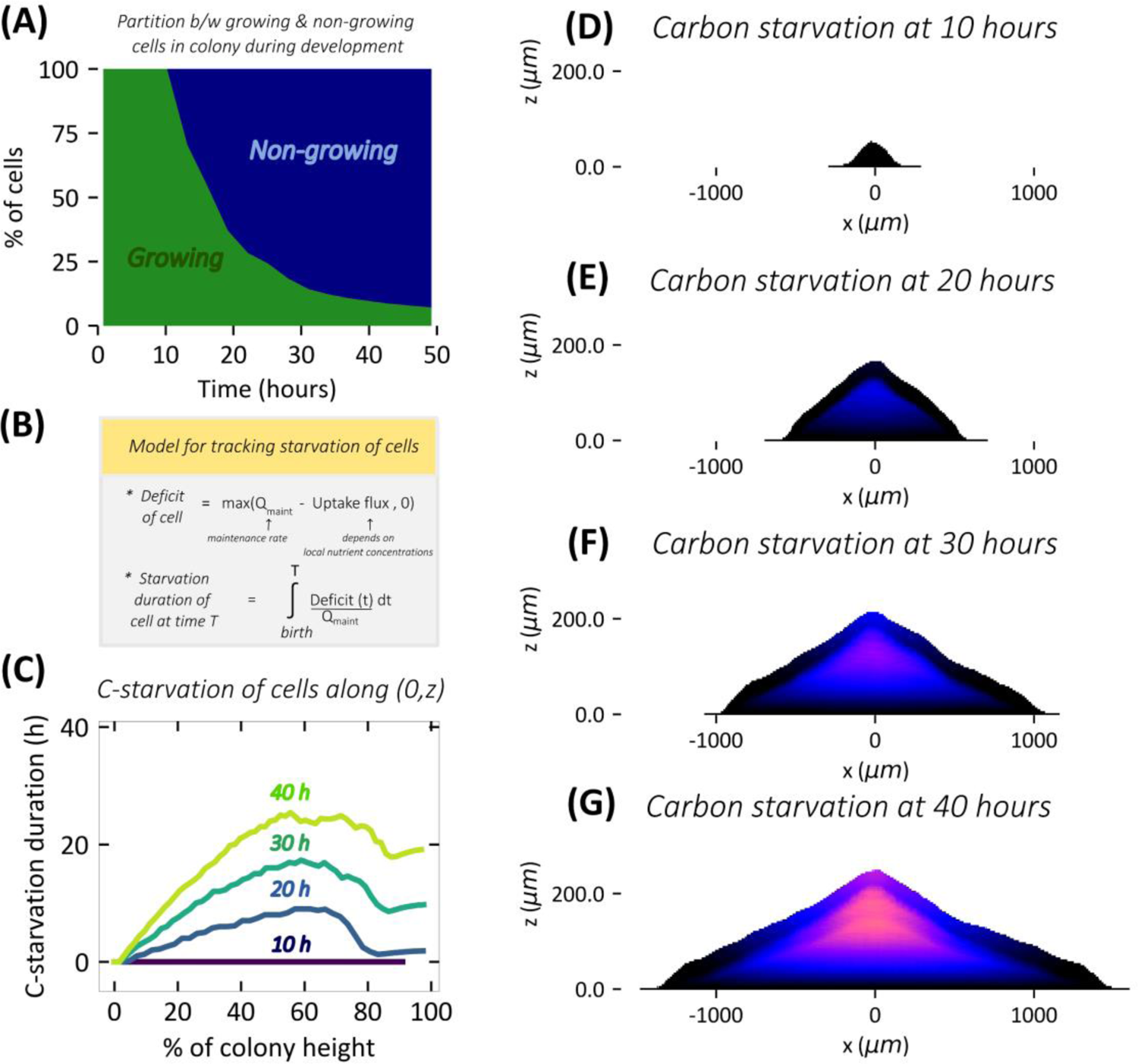
Progressive carbon starvation of cells in the interior during colony development. **(A)** The percentage of growing (green) and non-growing cells (blue) within the colony is plotted against the time of simulalted colony development. A cell is classified as growing if its instantaneous growth rate is greater than 0.01 h^-1^ and as non-growing otherwise. **(B)** A simplified description of our model to track the duration of nutrient starvation for cells within the colony (Further details are provided in **SI Appendix Section 2**). The starvation duration of cells (coded by color) within the colony at **(D)** 10 h, **(E)** 20 h, **(F)** 30 h, and **(G)** 40 h of simulated colony development. **(E)** The starvation duration of cells along the central vertical axis (0,z) of the simulated colony plotted against the vertical position from colony bottom (represented by % of the colony height) at 10 h, 20 h, 30 h and 40 h stages (coded by color) of colony development. Results here are for simulations with 20 mM initial glucose concentration in agar.

Given that the majority of the colony is non-growing beyond 20 hours (**Fig. 6A**), we sought to understand the spatiotemporal dynamics of nutrient starvation within the colony. Here we define starvation as a state in which the carbon uptake flux of a cell is less than the maintenance flux (**Fig. 6B**, see **SI Appendix Section 2** for further details). Our agent-based model enables us to track the starvation duration for each non-growing cell (**SI Appendix Section 2**). **Fig. 6D-G and Supplementary Video 9** provide a visual representation of the spatiotemporal dynamics of starvation within the colony. Virtually no starvation is present at 10 h while after that a starving central region emerges and expands in size. Upon plotting the starvation duration of cells in the center as a function of colony height (**Fig 6C**), we find that it is the interior region of the colony, in-between agar, and the colony top surface, that accumulates the longest duration of starvation. This is maintained throughout the colony development after the onset of starvation, with the duration starvation increasing as the colony ages (**Fig 6C**).

### Starvation induces cell death in the colony interior

The inevitable physiological consequence of prolonged starvation is cell death (31). Given that a significant portion of cells in the colony interior are under starvation, we sought to characterize the extent of cell death within colony. We first quantified the relationship between the duration of starvation and probability of cell death by tracking the viability of *E. coli* cells subjected to a controlled duration of carbon starvation using batch culture in aerobic and anaerobic conditions (see Methods). In **Fig. 7A** we report the temporal dynamics of the optical density (OD_600_) of liquid batch culture of EQ59 strain subjected to glucose depletion at the indicated time (“time zero” in **Fig. 7A)**. **Fig. 7B** shows that the viability of culture drops exponentially at a much faster rate for anaerobic cultures (∼ 2 per day) in comparison to aerobic cultures (∼0.2 per day). From this empirical relationship between the duration of starvation and death, our agent-based model is able to predict the fraction of dead cells within a colony based the duration of starvation and whether the cell experiences an aerobic or an anaerobic environment (see **SI Appendix Section 2D** for details). The spatiotemporal dynamics of the predicted death zone during the first 50 hours of simulated colony development is reported in **Supplementary Video 10**. Snapshots reported in **Fig. 7D** and **Fig. 7E** show the death zone within a 16-hour old and a 38-hour old simulated colony and it is seen that death is predicted to be localized in the interior anoxic core of the colony (**Extended Data Fig. 7AB**).

**Fig. 7.**
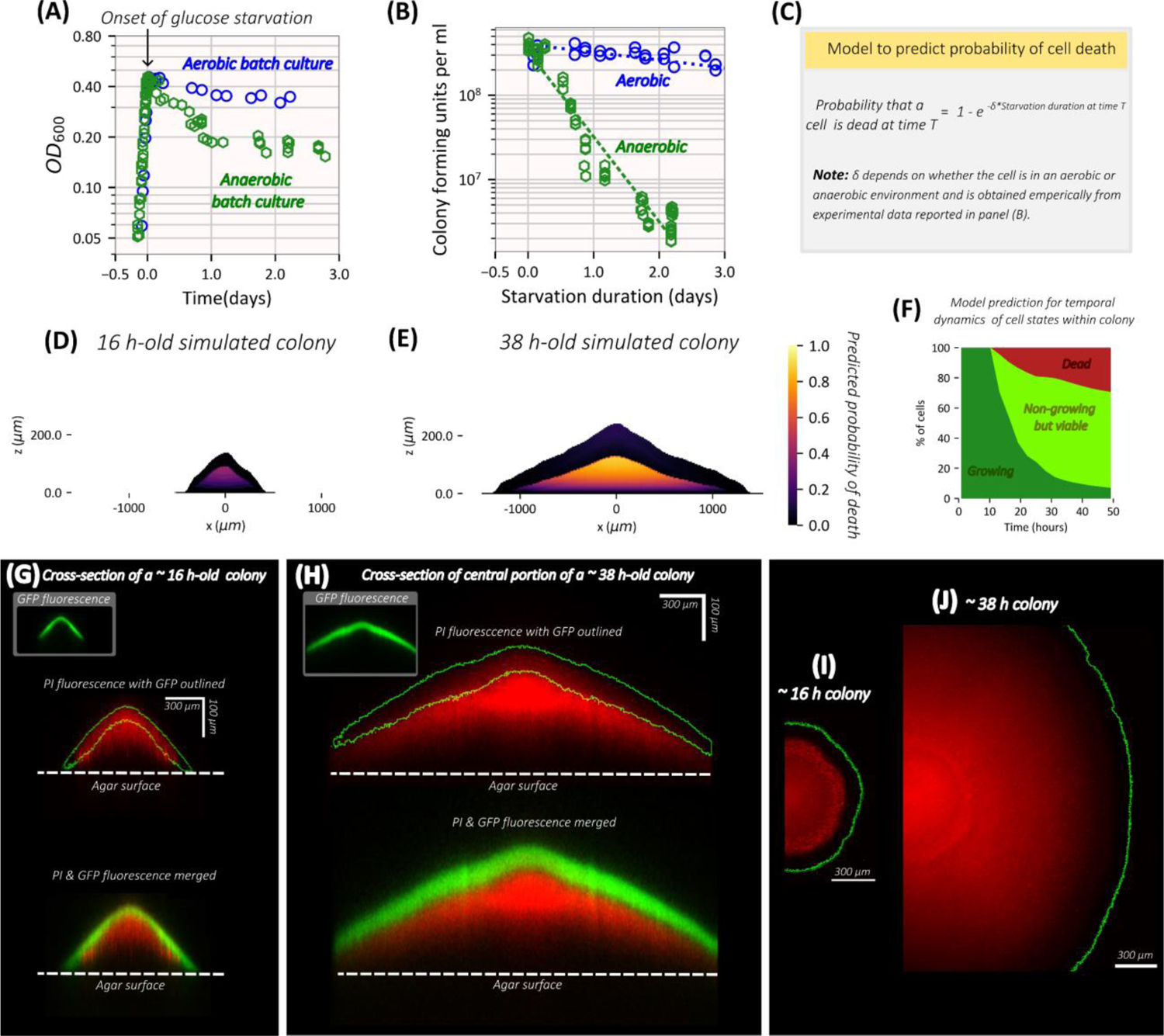
Prolonged starvation causes cell death in colony interior within 2 days of development. **(A)** Temporal dynamics of optical density measured at 600 nm (OD600) of aerobic (in blue) and anaerobic batch cultures (in green) that are under steady state exponential growth until ∼ 0.4 OD at which point glucose in the medium runs out and culture enters glucose starvation. **(B)** The number of viable cells per ml (colony forming units / ml) of culture is plotted against the duration of glucose starvation (days) for the aerobic cultures (in blue) and anaerobic cultures (in green). By fitting data to a curve of the form *a*e^−δt^, the death rate δ for aerobic and anaerobic glucose starvation are found to be 0.21 per day and 2.32 per day respectively. The blue and green dashed lines represent the fitted curve for aerobic and anaerobic glucose starvation respectively. **(C)** Illustration of model to predict the death probability of a cell within the colony based on the duration of starvation introduced in **Fig. 6B**. Model prediction for the probability of death (coded by color) averaged within a 4 μm x 4 μm grid at each spatial location within a **(D)** ∼ 16 h-old simulated colony and **(E)** ∼ 38 h-old simulated colony. Simulation results presented in **(D)** and **(E)** are with 20 mM initial glucose concentration. **(F)** Temporal dynamics of different cell states: growing cells (dark green), non-growing but viable cells (light green) and dead cells (red) within simulated colony. Propidium iodide fluorescence (in red) in an optical cross-section at the center of a **(G)** ∼ 16 h old colony and **(H)** a ∼ 38 h-old colony obtained using two-photon microscopy (see Methods). The fluorescence from GFP consitutively expressed by cells (see inset) is indicated by green outlines in top row of **(G)** and **(H)** and the fluorescence from GFP (green) and PI (red) merged together are shown in bottom row of **(G)** and **(H)**. Fluorescence along off-center cross-sections of a 38 h-old colony are reported in **Extended Data Fig. 8**, Maximum intensity z-projection of propidium iodide fluorescece (in red) near the peripheral edge of a **(I)** ∼ 16 h and **(J)** ∼ 38 h-old colony with the edge of the colony determined by GFP fluorescence outlined in green. Colonies were grown on 1.5 % agar plates with 20 mM glucose, 10 mM ammonium chloride, 112 mM phosphate buffered saline and 2.5 μM of propidium iodide.

To gather experimental evidence to test our model prediction, we use propidium iodide fluorescence along the cross-sectional profile of a colony to identify the *death zone* within a colony (see Methods). Propidium iodide is a fluorescent cell-membrane impermeant dye whose fluorescence intensity increases approximately 20 to 30-fold upon binding to DNA and is used to detect dead cells with compromised cell membranes (31, 33). In order to image fluorescence emitted by cells within the bulk of an intact colony which is a highly dense object, we use two-photon microscopy with high light penetration capabilities (34, 35) (See Methods). In **Fig. 7G and 7H** we report the fluorescence signal from the red emission wavelength band corresponding to propidium iodide fluorescence along the central cross-section of a 16-hour and a 38-hour old colony correspondingly with the outline of GFP fluorescence intensity indicated in green. It is indeed found that the intensity of propidium iodide fluorescence peaks at around the mid-point of colony height for 16 h and 38 h old colonies (**Fig. 7GH and Extended Data Fig. 7CD**) and this is in qualitative agreement with our model predictions of the spatial location of the *death zone* within the colony (**Fig. 7DE and Extended Data Fig. 7AB**).

## DISCUSSION

In this work, we study the development of non-EPS-producing *E. coli* colonies growing on hard agar. After an initial transient period of exponential growth, the colony expands at a constant radial speed for two days. On the other hand, the vertical expansion slows down substantially beyond the establishment phase (the initial 20-25 hours). To understand the differential behavior of radial and vertical expansion beyond the establishment phase, we have extended a previously introduced hybrid agent-based/continuum model describing the mechanical interactions of individual cells within colony and the diffusion and metabolism of key nutrients. To accommodate the large computational demand of an agent-based model beyond the establishment phase, we adopted a (1+1)-dimensional model focusing on a cross-section of the colony. Complex metabolic interactions in different regions of the colony are captured by including three key modes of metabolism: aerobic growth on glucose, anaerobic growth on glucose, and aerobic growth on acetate, along with cellular maintenance (both aerobic and anaerobic) in the absence of growth. Our agent-based model with a unique implementation of cell-level surface tension, described originally in Warren et al (23), was developed by incorporating commonly adopted modelling approaches (19, 36–44) to simulate the mechanical interactions within dense bacterial populations. As a novel addition in this study, we couple this agent-based model with a realistic metabolic model including anaerobic metabolism and maintenance, with parameters fixed by quantitative measurements. This model is further expanded to predict death probability for starving cells using experimentally measured death rates. Finally, by tracking both the movement and metabolic state of individual cells over time, we are able to obtain a spatiotemporal picture of progression of starvation and death within the colony.

One of the earliest studies of colony expansion by Pirt (12) hypothesized a depletion of nutrient concentration in the radial direction from the periphery into the colony to account for the linear radial expansion observed. This hypothesis is commonly adopted by the research community and motivates the use of, e.g., Fisher equation to describe radial colony expansion (45, 46). Warren et al (23) instead indicated the limitation of radial colony expansion to be mechanical factors that limited the width of radial expansion zone (the peripheral monolayer of cells) by forcing buckling and consequent verticalization (4, 36, 42, 47) of interior cells. However, those results were put forth for the establishment phase when oxygen penetration was not expected to be an issue. Simulations performed in this study with the more comprehensive metabolic model in fact found a sharp drop of oxygen concentration from the periphery into the colony beyond the establishment phase (**Extended Data Fig. 5D**). This finding potentially supports Pirt’s hypothesis (1), with oxygen being the limiting nutrient. On the other hand, the presence or absence of oxygen makes only a minor difference to the growth rate of *E. coli* on glucose (**SI Appendix Table 1**). We thus scrutinized the peripheral region more closely: Our results show that both oxygen and glucose are well above their corresponding K_c_ values at the buckling width for the colony throughout the first 50 h of colony development (**Extended Data Fig. 9B**). Thus, neither oxygen nor glucose was depleted in the region where radial expansion takes place; instead, the width of the radial expansion zone was set by mechanical instability that forced verticalization as was found in the establishment phase. This picture on radial expansion is summarized in **Fig. 8A**.

**Fig. 8.**
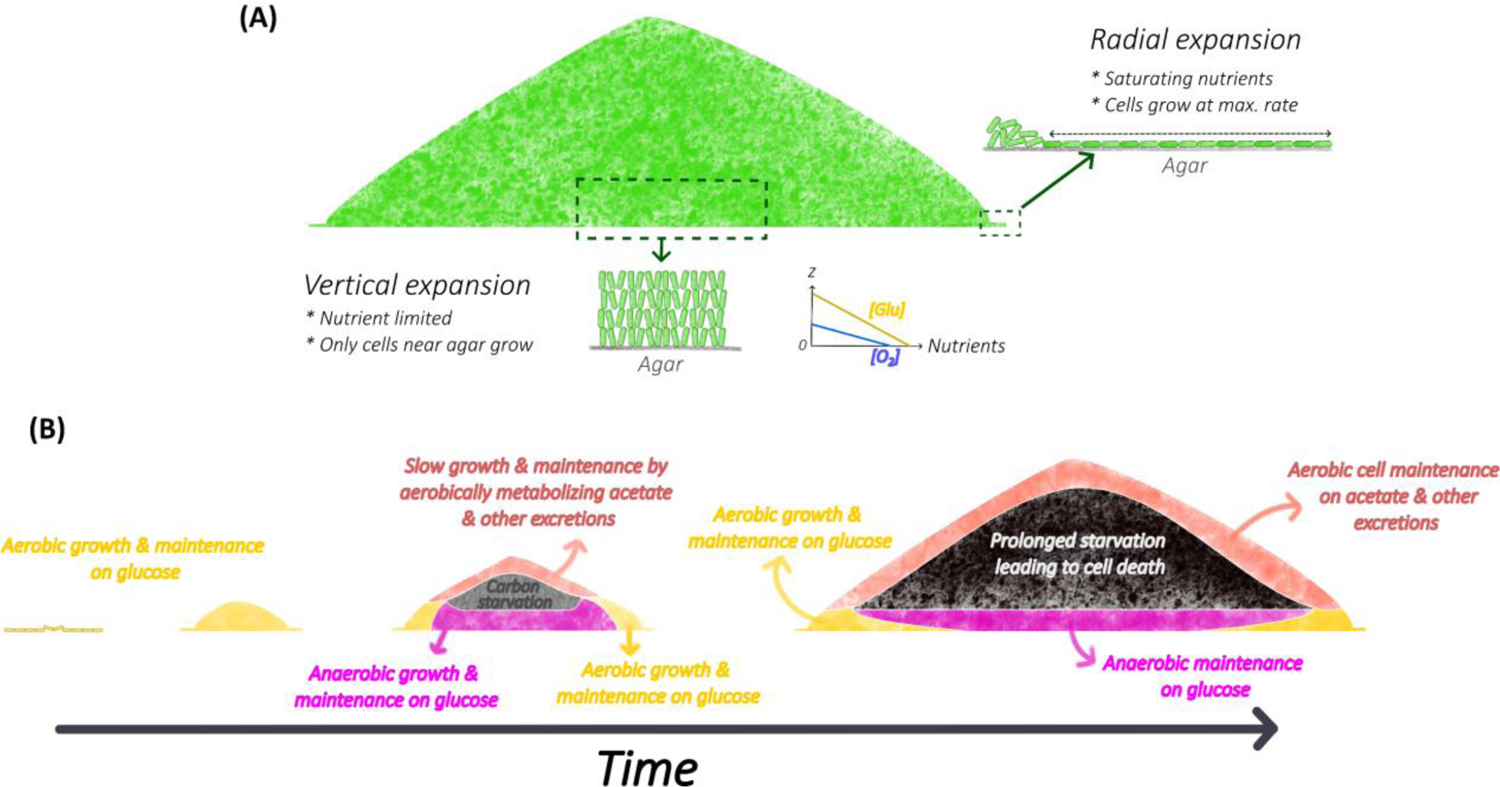
The developmental program of colony expansion dictated by mechanical constraints and emergent nutrient dynamics. **(A) *Overview of the mechanism for colony expansion:*** Radial expansion is determined by the growth of the peripheral monolayer of horizontally oriented cells which experience a saturating concentration of nutrients (glucose and oxygen) and grow at the maximum rate. Vertical expasnion is determined by the growth of vertically oriented interior cells located near the colony-agar interface which is nutrient limited. **(B) *Overview of the spatiotemporal dynamics of metabolism within colony:*** Initial stages of colony development involve the transition from a monolayer to a 3D colony during which aerobic glucose metabolism dominates. Later, due to oxygen depletion, cell growth from anaerobic glucose metabolism dominates over cell growth from aerobic glucose metabolism which is restricted only to the peripheral edges. Due to high consumption flux, glucose is limited to a few μm away from the agar interface. Further, excretions from anaerobic metabolism such as aceate contribute to slow growth and maintenance at the colony-air interface. The colony interior experiences starvation since acetate cannot be metabolized without oxygen which is restricted to a few μm from the colony-air interface. Thus, as time progresses, due to glucose depletion, growth (by aerobic glucose metabolism) is restricted to peripheral edges of the colony with cell maintenance from anaerobic glucose metabolism near agar interface and cell maintenance from aerobic acetate (& other excretions) metabolism at the air interface. Further, due to prolonged starvation, a zone of dead cells form in the colony interior.

Our simulations do find the sharp depletion of glucose inside the colony to be the primary cause of the vertical slowdown. This severe glucose depletion arises from the high cell density in the colony and is further accelerated by the high glucose consumption flux during anaerobiosis (**SI Appendix Table 1**). In recent years, there has been a renewed interest in understanding the vertical dynamics of biofilm formation on hard substrates. A comprehensive experimental study by Bravo et al (16) characterized the vertical dynamics of biofilm expansion for various microbial species with nanometer precision measurements of biofilm height. Their findings indicate that the vertical expansion kinetics of monoclonal biofilms formed by diverse microorganisms universally experience a slowdown beyond an initial fast expansion regime. Bravo et al explained this phenomenon based on a heuristic interface model wherein cell growth was assumed to be restricted to a thin layer at the colony-substrate interface. This assumption is however at odds with findings of other recent studies (19, 20, 48) which expected the excretion of fermentation products such as acetate also to play important roles at the colony-air interface. This apparent conflict is resolved in our study by recognizing that the contribution of the excreted acetate to colony growth is minimal at a quantitative level (**Supplementary Video 7**). This finding stems from the low maximum growth rate (∼0.4/h) of *E. coli* on acetate, coupled with the fact that a high concentration of acetate (∼5 mM) is required for cells to grow at half of the maximum rate (**Extended Data Fig. 10**). Our simulations find acetate to accumulate to ∼8 mM in the colony early on (< 20 h), dropping to well below 5 mM after that (**Extended Data Fig. 6A**), owing to a combination of effects including the consumption of acetate by cells, reduced excretion of acetate due to glucose depletion, and diffusion of acetate from colony into the agar. These findings on vertical colony expansion are summarized graphically in **Fig. 8AB**.

We regard the most significant finding of this work to be connecting the slowdown in vertical colony expansion to cell death, the inevitable physiological consequence of prolonged carbon starvation. Our model predicts a localized death zone within the colony interior away from the top and bottom interfaces within 1-2 days of colony development (**Fig. 7DE**), validated by observing propidium iodide fluorescence along the cross-section of intact colonies using two-photon microscopy (**Fig. 7G-J and Extended Data Fig. 8**). In our simulations, we find that acetate metabolism plays a key role in cell maintenance at the colony-air interface (where oxygen can be utilized). Thus, even though acetate contributes minimally to cell growth (**Supplementary Video 7**), it is a key factor keeping the top layers of the colony viable by contributing to cell maintenance (**Supplementary Video 8**). Factors governing cell viability/death are summarized in **Fig. 8B**.

In our metabolic model, we only included acetate as the fermentation product due to its large excretion flux (49). There are of course other excretion products that can contribute to cell metabolism close to the colony surface where oxygen is available. In this vein, a recent study by Diaz-Pascual et al (13) discussed the consumption of alanine by cells at the colony-air interface based on their spatially resolved transcriptomic data for *E. coli* colonies. Alanine can be utilized by cells as both a carbon and nitrogen source. In our experiments, changing the concentration of ammonium did not affect the vertical expansion of colony (**Extended Data Fig. 3**), suggesting that the nitrogen source is not the rate-limiting nutrient for colony growth. Further, since the excretion of alanine is much lower than acetate during fermentation (50), we expect alanine utilization as a carbon source to be lower than acetate utilization within the colony. Spatial transcriptomics done in Diaz-Pascual et al (13) did not reveal signatures of acetate crossfeeding by looking at the fold change in expression of genes such as ackA, pta and eutD. At low acetate concentrations (< 10 mM), it is likely that the ackA-pta pathway does not participate in acetate assimilation and instead the AMP-Acs pathway is expected to be primarily used by *E. coli* for acetate utilization (51). Further, genes involved in acetate metabolism (e.g. acs, aceA, aceB, etc.) are part of a larger group of carbon catabolic genes (52) which have an obligatory upregulation during carbon limitation (52–54). This upregulation of genes involved in acetate metabolism during carbon limitation occurs regardless of whether acetate is utilized or not. Given that the bulk of the colony is expected to be under carbon limitation (**Supplementary Video S8,9**), it would be difficult to detect signatures for acetate crossfeeding by looking at fold-change in expression of genes involved in acetate metabolism. This is because the only cells which are expected to not express acetate metabolism genes are the cells growing on glucose which incidentally form a tiny fraction of the colony even as early as 12 hours into development (**Supplementary Video S7**). As seen from **Supplementary Video S8** (bottom row) and **Extended Data Fig. 6**, acetate utilization is predicted near the colony-air surface, which is in agreement with an earlier study by Cole et al. (19). Further, it should be noted that our observations do not rule out alanine uptake at the oxic region near the colony-air interface. Rather, it is unlikely that cells at the colony-air interface rely primarily on alanine for carbon and nitrogen since acetate (carbon source) and ammonium (nitrogen source) are both expected to be present at higher levels than alanine. Instead, the uptake of alanine at the colony-air interface could play a secondary role with acetate and ammonium being the primary carbon and nitrogen sources respectively.

In this work, we have not included the recycling of lysis product (e.g., amino acids and nucleotides) released by dead cells except indirectly through the adoption of an exponential death rate which is quantitatively influenced by the extent of such nutrient recycling (31, 55). Such lysis products could in principle fuel cell growth and maintenance at the colony-air interface where oxygen is available; however, the metabolic processes involved are much more complicated and will be the subject of a future study. We also note that high levels of acetate accumulation can in principle cause a toxic effect on cells within the colony (49, 56–58). In this study, we use a high concentration of phosphate buffer (∼112 mM) much exceeding the excreted acetate concentration (< 10 mM) to maintain a neutral pH throughout the colony. This may not generally be the case for colony growth in other conditions, and the effects of acetate-mediated toxicity on colony growth and cell viability will be discussed elsewhere.

What this study has uncovered through a combination of modelling and experiments are basic characteristics of growing bacterial colonies stripped off of the features of EPS-formation and cell differentiation that make the study of biofilms more difficult (2–4, 59). But the findings obtained here on the simplest non-EPS-producing colonies already reveal a complex pattern of spatiotemporal dynamics that occur presumably in all biofilms since the metabolic processes involved (aerobic and anaerobic growth on glucose, aerobic growth on fermentation products, aerobic and anaerobic maintenance of cell viability) are basic processes that govern all heterotrophic bacteria (60, 61). We thus believe the approaches and findings here represent important first steps towards a systemic quantitative understanding of more complex biofilms that also involve EPS formation and cell differentiation (59, 62). Indeed, colony-forming microorganisms can exploit the obligatory nutrient gradients formed at different locations and time in developing colonies as spatiotemporal cues to signal the development of more complex spatial structures and temporal growth patterns (11, 63–68).

## METHODS

### Experimental methods

#### Bacterial strains

The strains, EQ54 and EQ59 of *E. coli* K12 used in this study are derived from NCM3722 (69). EQ54 is constructed by the deletion of motA gene and EQ59, in addition to a deletion in motA gene harbors constitutive expression of green fluorescent protein (GFP). The details of EQ54 strain construction are described in Ref. (70) and EQ59 strain construction is described in Warren et al. 2019 (23).

#### Growth medium

For studying colony growth kinetics minimal medium agar plates were used. The composition of each agar plate comprises of 1.5% agar (w/v) along with a defined concentration of glucose (carbon source), 10 mM NH_4_Cl (nitrogen source), 42 mM NaCl, 0.4 mM MgSO_4_.7H_2_0, 5.5 mM K_2_SO_4_ along with phosphate buffer to maintain a neutral pH (∼ 7.2) comprising of 77.5 mM K_2_HPO_4_ and 34.5 mM KH_2_PO_4_. The total media volume used for each petri-dish is ∼ 16 mL. For experiments to image the death zone within colonies, 2.5 μM of propidium iodide was added to the above growth medium during preparation of minimal media agar plates.

#### Batch culture growth

Batch culture growth is performed in a 37°C water bath shaker (220 rpm). Cells from a glycerol stock stored at −80°C are streaked on an LB agar plate and incubated overnight at 37°C. A single colony from this plate is picked and inoculated into LB broth and grown in a shaking water bath for several hours at 37°C as seed cultures. Seed cultures are then transferred into minimal medium and grown overnight at 37°C in a shaking water bath as pre-cultures. For batch culture measurements, overnight pre-cultures are diluted to around 0.01 to 0.02 OD_600_ in fresh minimal medium and grown at 37°C in a shaking water bath as experimental cultures. For batch culture growth in minimal medium, the same medium composition as the minimal medium for colony growth excluding the 1.5% agar was used.

For anaerobic batch cultures, the medium used is the same as used for aerobic cultures. To maintain an anerobic environment for cell growth, Hungate tubes (16 mm x 125 mm) with 7 mL of the medium are shaken at 270 rpm under 7 % CO_2_, 93 % N_2_ atmosphere pressurized to 1.5 atm for 75 minutes prior to being used for experiments. Cultures were transferred into and out of Hungate tubes with disposable sterile syringes.

#### Colony growth in minimal media agar plates

Seed cultures and pre-cultures were prepared as described above. Overnight pre-cultures are then diluted to OD_600_ around 0.01 to 0.02, in fresh minimal medium and grown at 37°C in a shaking water bath as experimental cultures. When the experimental culture reaches ∼ 0.2 OD, an aliquot of experimental culture is taken and diluted 10^-6^ fold in phosphate buffer and ∼ 100 ul of the diluted sample is plated on to minimal medium agar plates. The liquid sample is then spread with sterile glass-beads to yield ∼ 10 colony forming units on the agar plate. The plates are dried under a flame for roughly 10 minutes, wrapped with parafilm and kept in an inverted position in a 37°C incubator. At periodic intervals the agar plates are taken out of the incubator and unwrapped for microscopy measurements which last roughly ∼5 minutes. Post-microscopy the plates are wrapped with parafilm and placed in an inverted position in a 37°C incubator again.

#### Batch culture viability measurements

To measure viability of batch cultures under glucose starvation, seed cultures and pre-cultures were prepared as described above. Overnight pre-cultures are then diluted to OD_600_ around 0.01 to 0.02, in fresh minimal medium with a limiting concentration of glucose such that at ∼ 0.4 OD_600_, glucose runs out in the medium and cells in the culture enter stationary phase. The cultures are maintained at 37°C grown in a shaking water bath throughout the experiment. To measure the viability of cells in the glucose-starved culture, an aliquot of culture is sampled at periodic intervals post-glucose runout, diluted, and plated on LB agar plates. The liquid sample is then evenly spread on the agar plate with ∼ 5-6 sterile glass-beads. The number of colony forming units formed per mL of culture is taken to be a measure of the viability of cells in the culture. Only plates with colony forming units between ∼30-300 c.f.u. on the agar plate were counted (manually) to obtain c.f.u. per mL of culture.

#### Confocal microscopy

Colonies growing on agar plates are imaged using a Leica TCS SP8 inverted confocal microscope equipped with a stage housed in a 37 ° C temperature-controlled box. The agar plates are uncovered and placed in an inverted position on the stage and a xyz-stack scan is performed to obtain a 3D image of the colony. Fluorescence from GFP that is constitutively expressed by cells is detected using excitation by a 488 nm diode laser and detected with a 10x/0.3 NA air objective and a high sensitivity HyD SP GaAsP detector. A montage of tile-scans are created and stitched together to form a single 3D image using a custom python script.

#### Image processing and analysis

The cross-sectional profile of a colony is obtained from the stitched z-stack confocal images by computing the bounding radius for each z-coordinate. From the cross-sectional profile, the maximum radial dimension which defines the colony bottom is reported as the radius and the height is computed as the maximal distance from the bottom at which the bounding radius is still non-zero. Further, for estimating the colony volume, the colony is treated as a stack of thin disks i.e., for each z-step the colony is approximated as a disk of thickness corresponding to the z-step (∼ 5 μm) and a radius corresponding to the radial dimension at the particular z-coordinate. Then, the colony volume is estimated as the sum of the volume of these thin disks that make up the colony. The above computations were performed using a custom python script.

#### Two-photon microscopy

Two-photon laser scanning microscopy with adaptive optics (AO-TPLSM) (35) was employed to image the colonies either ∼16 hours or ∼38 hours after their growth on agar plates. AO-TPLSM was performed with a 10X air objective (0.5 NA; Thorlabs, TL10X-2P). The system aberration was corrected by a deformable mirror (ALPAO, DM97-15), as described in Ref. (34), for better image quality. Propidium iodide fluorescence was excited by a tunable femtosecond laser (Coherent, Chameleon Discovery) tuned to 1070 nm, filtered through a red bandpass filter (Semrock, FF01-593/46-25), and detected with a silicon photomultiplier (Hamamatsu, C13366-3050GA). The fluorescence from GFP in the same colony was excited at 930 nm wavelength, filtered through a green bandpass filter (Semrock, FF01-530/55-25), and detected with the silicon photomultiplier. The post-objective power was approximately 50 mW for imaging the propidium iodide fluorescence and 20 mW for GFP fluorescence imaging.

### Computational model

Our model is comprised of three components: an agent-based model for individual cells; a metabolic model to determine local cell growth rate; and a system of reaction-diffusion partial differential equations (PDEs) for modeling the spatiotemporal dynamics of metabolism inside the colony. The agent-based model for individual cell activities was constructed in our previous work, Warren et al. 2019 (23) and is reviewed in **SI Appendix Section 1**. Here, we present a brief description of the model to determine local cell growth rate and the reaction-diffusion equations for spatiotemporal dynamics of metabolism with additional details provided in **SI Appendix Section 2**. We also provide a brief description of the numerical algorithm used to simulate colony growth with additional details provided in **SI Appendix Section 3**.

### Growth modes and local cell growth rate

We include three modes of cell growth in our model: aerobic growth on glucose, anaerobic growth on glucose, and aerobic growth on acetate (**Fig. 2A-C**). The local cell biomass growth rate λ = λ^(⃗^r⃗, *t*) at a spatial point *r^→^* within the colony and time *t*, the main quantity that connects the continuum and discrete parts of our hybrid model, is a weighted sum of the growth rates corresponding to the three different growth modes, given by

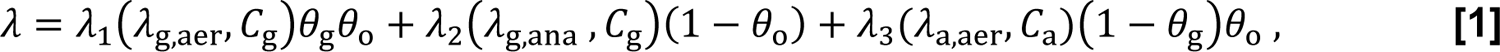

where λ_1_, λ_2_ and λ_3_represent the local growth rates for aerobic growth on glucose, anaerobic growth on glucose, and aerobic growth on acetate, respectively. These local growth rates depend on the local concentrations of glucose *C*_g_ = *C*_g_(r^→^, *t*), oxygen *C*_o_ = *C*_o_^(⃗^r⃗, *t*), and acetate *C*_a_ = *C*_a_(r^→^, *t*), and also on the maximum growth rates, λ_g,aer_, λ_g,ana_, and λ_a,aer_, corresponding to each of the three growth modes, respectively. Further, θ_g_, θ_o_, θ_a_ represent Monod kinetic forms, namely,

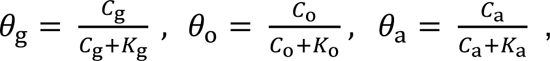

where *K*_g_, *K*_o_, and *K*_a_ are the Monod constants for glucose, oxygen, and acetate, respectively. The factors θ_o_ and (1− θ_o_) represent the weights assigned for aerobic growth and anaerobic growth, respectively, and the factors θ_g_ and 1 − θ_g_are the weights for the growth from glucose and acetate, respectively.

In a model which does not account for cell maintenance, the local rates λ_1_, λ_2_ and λ_3_ for different growth modes would traditionally be defined using Monod kinetic forms.

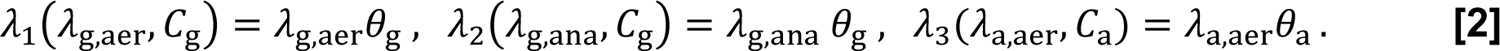

However, to incorporate the effects of cell maintenance on the local cell growth rate λ (Eq. **[1]**) we appropriately modify the local growth rates λ_1_(λ_g,aer_, *C*_g_), λ_2_(λ_g,ana_, *C*_g_), and λ_3_(λ_a,aer_, *C*_a_) for cell maintenance which is described in **SI Appendix Section 2**. For instance, considering aerobic growth on glucose, we introduce a threshold value *C*^∗^ for glucose concentration *C*_g_ and modify the corresponding growth rate λ_1_(λ_g,aer_, *C*_g_) to be 0 if *C*_g_ ≤ *C*^∗^ and if *C* > *C*^∗^ we use an appropriately shifted version of the Monod curve to determine the growth rate. The growth rates corresponding to the other two modes, λ_2_(λ_g,ana_, *C*_g_) and λ_3_(λ_a,aer_, *C*_a_) are similarly modified. A detailed description of the determination of the threshold concentrations, as well as the explicit formulas for the local cell growth rates are provided in **SI Appendix Section 2**.

#### A continuum model for spatiotemporal dynamics of metabolite gradients

In our simulations, a defined initial concentration of glucose is set in the agar region. Glucose diffuses within the agar and into the colony region where it is taken up by cells for either growth or maintenance in the presence or absence of oxygen. During growth on glucose, cells excrete acetate which together with oxygen can also be taken up by cells for growth and maintenance. We describe these metabolic processes with a coarse-grained model through a set of reaction-diffusion PDEs together with appropriate initial and boundary conditions for the local concentrations *C*_g_, *C*_o_, and *C*_a_ of glucose, oxygen, and acetate respectively. Referring to **Fig. 2E**, the general form of the reaction-diffusion equations modeling the spatiotemporal dynamics of the metabolite concentrations *C* = *C*_g_, *C*_o_, or *C*_a_ is given by,

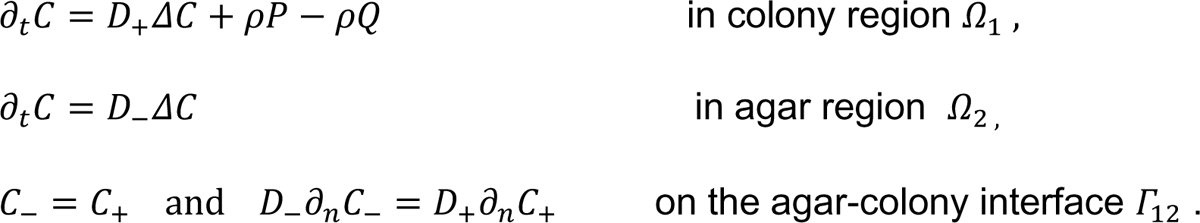

Here, *D*_−_ and *D*_+_ are the diffusion coefficients for the metabolite with concentration *C* in the agar and colony regions Ω_1_ and Ω_2_, respectively, *C*_−_ = *C* in agar and *C*_+_ = *C* in colony, and ∂_n_ denotes the normal derivative in the direction from agar to colony. If *C* = *C*_g_, *C*_o_, or *C*_a_, then we denote the diffusion coefficients by *D*_g,−_, *D*_g,+_, *D*_o,−_, *D*_o,+_, or *D*_a,−_, *D*_a,+_, respectively. In the reaction term ρ*P* − ρ*Q*, ρ is the local cell density and *P* and *Q* are the excretion and uptake rates, respectively, of the metabolites within the colony. We model these rates for glucose *C*_g_, oxygen *C*_o_ and acetate *C*_a_ as follows,

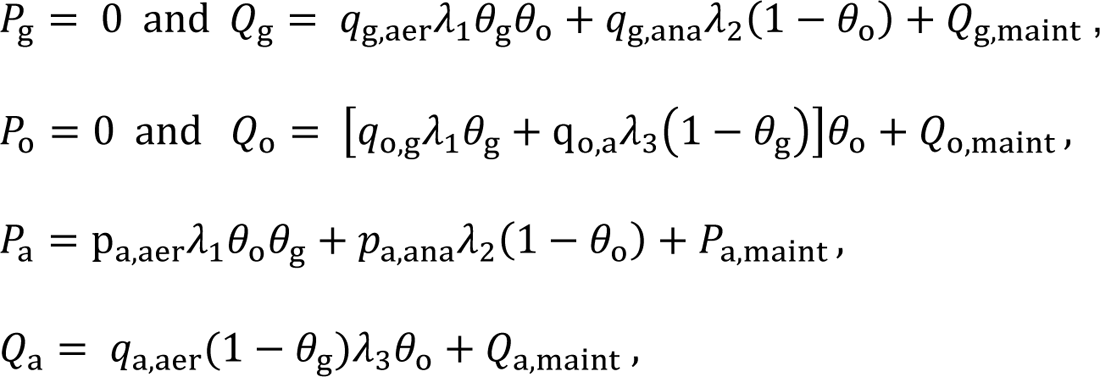

Here λ_1_ = λ_1_(λ_g,aer_, *C*_g_), λ_2_ = λ_2_(λ_g,ana_, *C*_g_)and λ_3_ = λ_3_(λ_a,aer_, *C*_a_) are given in Eq. **[2].** The subscripted *p* and q are constant parameters: *q*_g,aer_and *q*_g,ana_ are the specific uptake rates for glucose during the aerobic and anaerobic growth, respectively; *q*_o,g_ and *q*_o,a_ are the specific uptake rates for oxygen during the aerobic growth on glucose and on acetate, respectively; *p*_a,aer_and *p*_a,ana_are the specific excretion rates for acetate during aerobic and anaerobic growth on glucose, respectively; and *q*_a,aer_is the specific uptake rate for acetate during aerobic growth on acetate. Further, *Q*_g,maint_, *Q*_o,maint_and *Q*_a,maint_correspondingly represent the uptake rate of glucose, oxygen, and acetate for maintenance, while *P*_a,maint_represents the excretion rate of acetate arising from cell maintenance. Explicit mathematical expressions for the maintenance uptake and excretion rates of metabolites are provided in **SI Appendix Section 2** and the parameter values are listed in **SI Appendix Table 1**.

Flux-free boundary conditions are imposed for the concentrations of glucose *C*_g_and acetate *C*_a_ on the boundaries surrounding the colony and agar regions, i.e., ∂_*n*_*C*_g_ = 0 and ∂_*n*_*C*_a_ = 0 on Г_01_ ∪ Г_02_ ∪ Г_s_ ∪ Г_b_, (**Fig. 2E**). The boundary conditions for the concentration of oxygen are ∂_n_*C*_o_ = 0 on Г_s_ ∪ Г_b_ and *C*_o_ = *C*_o,0_ on Г_01_ ∪ Г_02_, where *C*_o,0_is a constant oxygen concentration. The initial glucose concentration is set to be a constant in the agar region, whereas the initial acetate concentration in agar is set to be 0.

### Numerical algorithm

Our simulations of colony growth are done iteratively with each iteration consisting of the following main steps: (1) Given the spatial coordinates of all the individual cells within the colony, we determine a coarse-grained, smoothened colony boundary. (2) With the colony boundary defined from the previous step, we solve the system of reaction-diffusion equations with appropriate boundary conditions in both the agar and colony regions to update the concentrations of glucose, oxygen, and acetate. (3) Given the local concentration of the metabolites from the previous step, we update the local growth rate for cells in the colony region. (4) Using the local cell growth rates computed in the previous step, we simulate the growth, division, and movement of all the individual cells within the colony with another nested loop of iteration with a smaller time step. Details of the boundary conditions, numerical methods for solving the set of reaction-diffusion equations (71, 72) and the numerical schemes (73) used to simulate the velocity and positions of individual cells can be found in **SI Appendix Section 3**.

## ACKNOWLEDGEMENTS

We thank Xiongfei Fu and Hwa lab members for helpful discussions. This work was supported in part by NIH through grant U24 EB028942 (DK) and by NSF through grants MCB-2029580 (PS), MCB-2029480 (JJD), DMR-1702321 (JJD), PHY153264 (DK), DMS-2208465 (BL) and MCB-2029574 (TH and BL).

## AUTHOR CONTRIBUTIONS

M.W. and T.H. conceived of the experimental study and M.W. performed initial investigations. H.K., M.W., T.C., and T.H. designed experiments and H.K., T.C., K.S., and B.R.T. performed experiments; P.S., M.W. and B.L. designed the numerical study and P.S. developed the parallelized simulation software; H.K., P.S. performed simulations with contributions from D.G. under the supervision of B.L.; H.K., M.W., M.M. and B.R.T. contributed to modelling with supervision from B.L, and T.H.. P.Y. performed two-photon microscopy imaging with the supervision of D.K.; T.C. designed image analysis pipeline for colony dimension measurements. H.K, J.D., B.L. and T.H. wrote the manuscript with input from all the authors.

## Supplementary Figures

**Extended Data Fig.1.**
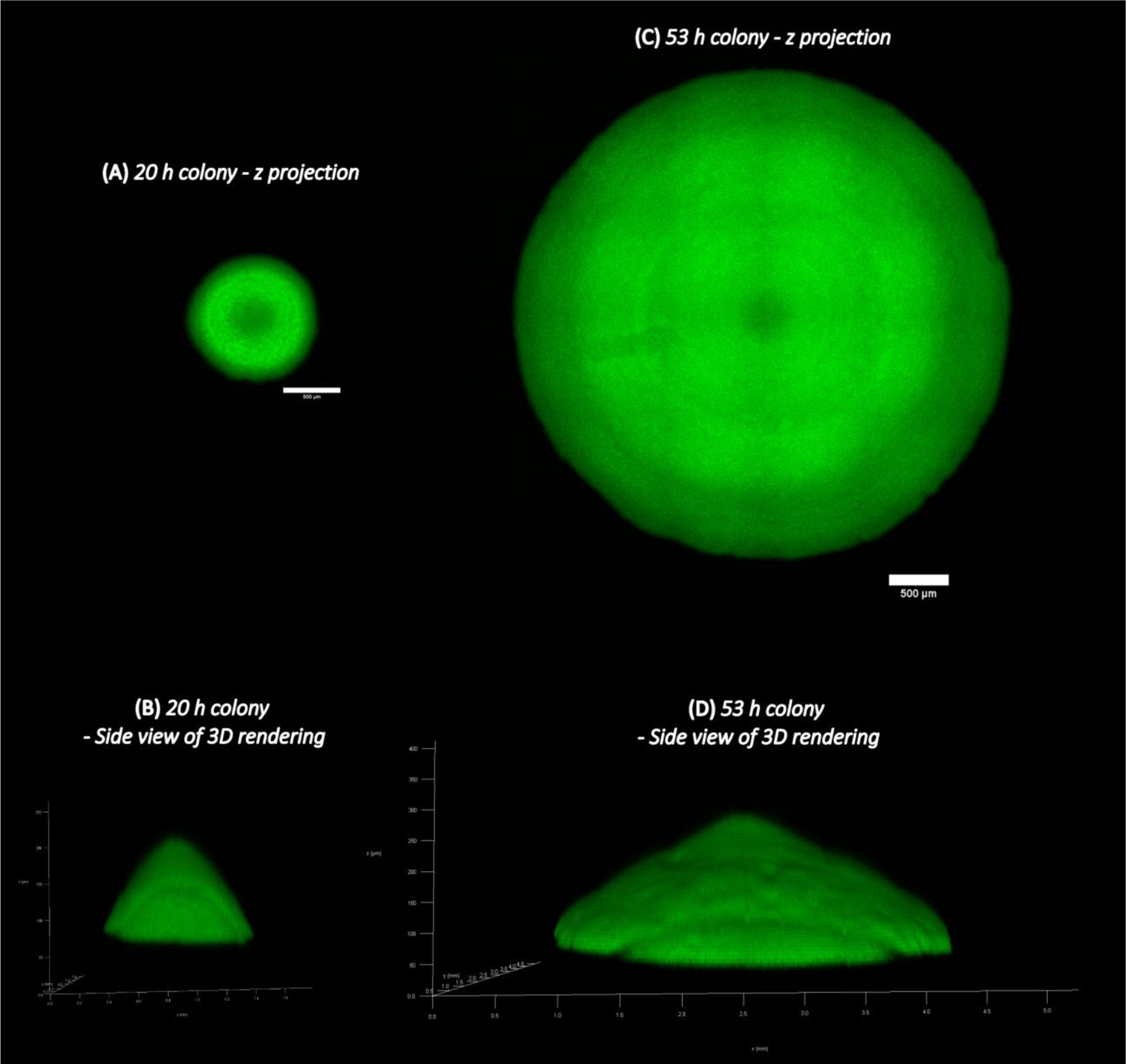
Confocal microscopy images of E. coli colonies. **(A)** z-projection and **(B)** side-view of 3D rendering from microscopy images of a ∼ 20 h old EQ59 *E. coli* colony. **(C)** z-projection and **(D)** side-view of 3D rendering from microscopy images of a ∼ 53 h old EQ59 colony. 1.5 %(w/v) agar plates prepared with 20 mM glucose, 10 mM ammonium chloride, and 112 mM phosphate buffered saline were used.

**Extended Data Fig.2.**
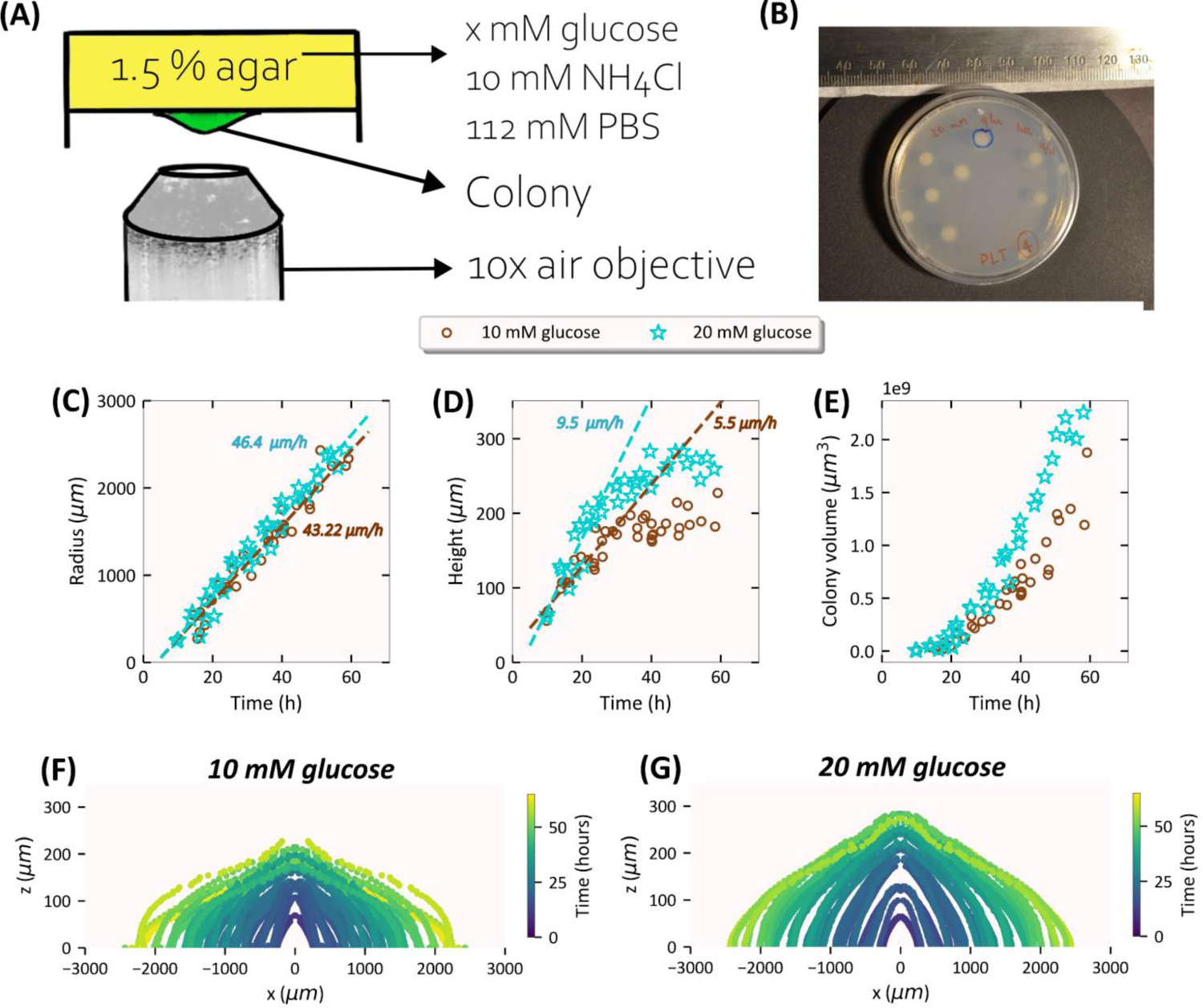
Overview of experimental set-up to measure colony dimensions and repeatability of observations. **(A)** An illustration of the experimental set up to measure the dimensions of a colony growing on an agar plate using an inverted confocal microscope. **(B)** A photograph of a typical 60 mm x 15 mm agar plate in our experiments with *E. coli* colonies at ∼ 2 days post-inoculation. The number of colonies per plate is kept low at roughly ∼ 10 colony forming units (c.f.u.). **(C)** The radius (μm), **(D)** The height (μm) and **(E)** The volume (μm^3^) of colonies from multiple experiments performed on different days plotted against the time (h) post-inoculation. Brown circles represent colonies grown on minimal media plates with 10 mM glucose while cyan stars represent colonies with 20 mM glucose as the carbon source. The cross-sectional profiles of colonies from multiple experiments performed on different days with **(F)** 10 mM glucose and **(G)** 20 mM glucose as carbon source shown for various times (coded by color) post-inoculation.

**Extended Data Fig.3.**
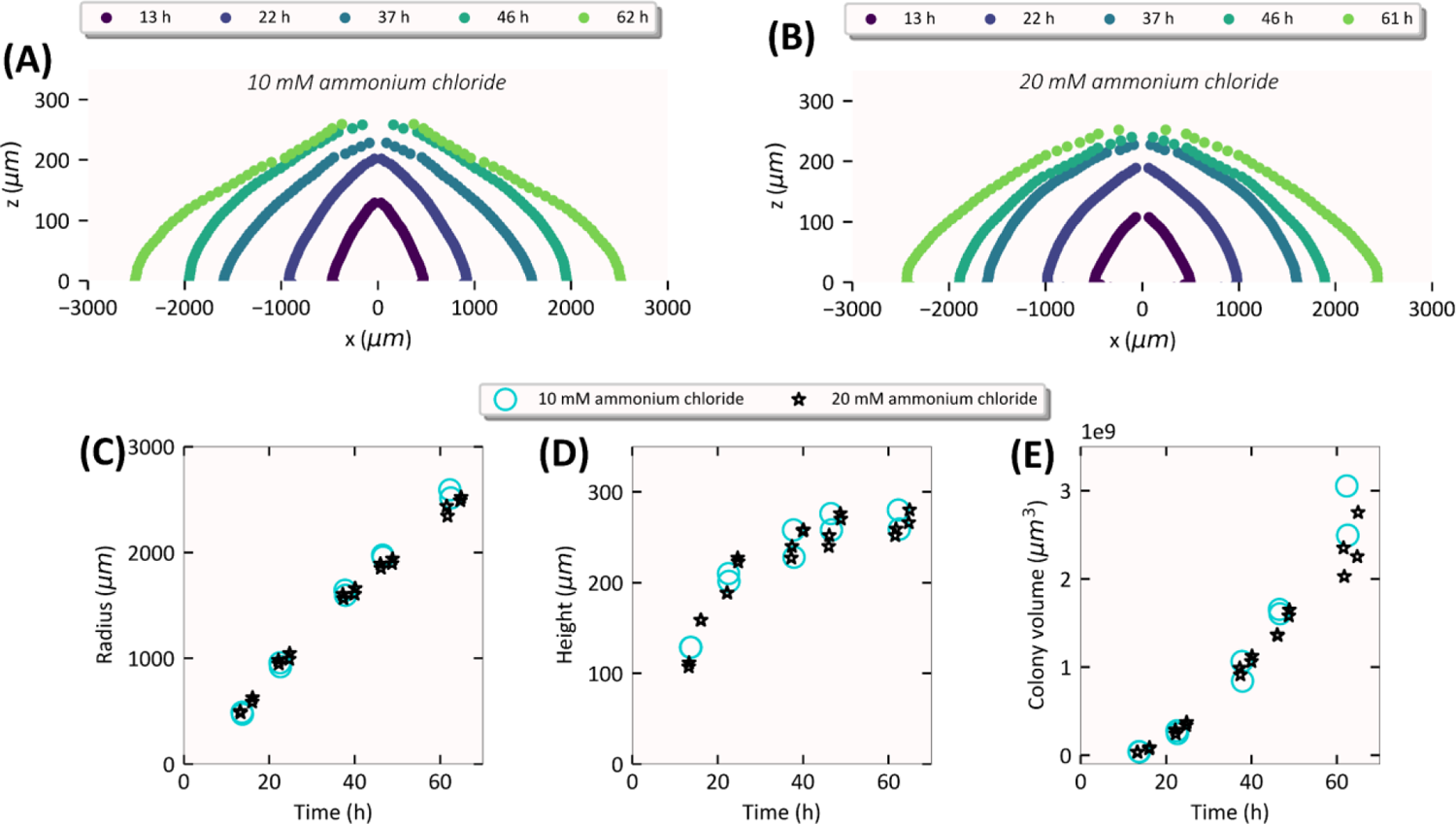
Colony expansion is not limited by the initial concentration of nitrogen source. Expansion kinetics of EQ59 *E. coli* colonies on 1.5 %(w/v) agar plates prepared with 20 mM glucose, a defined ammonium chloride concentration (10 mM, 20 mM), and 112 mM phosphate buffered saline (PBS) at various times post-inoculation as a single cell. The seeding density of colonies is such that there are ∼10 well-separated colony forming units on a petri dish which is 60 mm diameter and has ∼ 8 mm media depth with a total media volume of ∼ 16 ml. The cross-sectional profile of a colony grown on a minimal media hard agar plate with **(A)** 10 mM ammonium chloride and **(B)** 20 mM ammonium chloride as carbon source shown for various times (coded by color) post-inoculation. **(C)** The radius (μm), **(D)** The height (μm) and **(E)** The volume (μm^3^) of the colonies plotted against the time (h) post-inoculation. Cyan circles represent colonies grown on minimal media plates with 10 mM ammonium chloride while black stars represent colonies with 20 mM ammonium chloride.

**Extended Data Fig.4.**
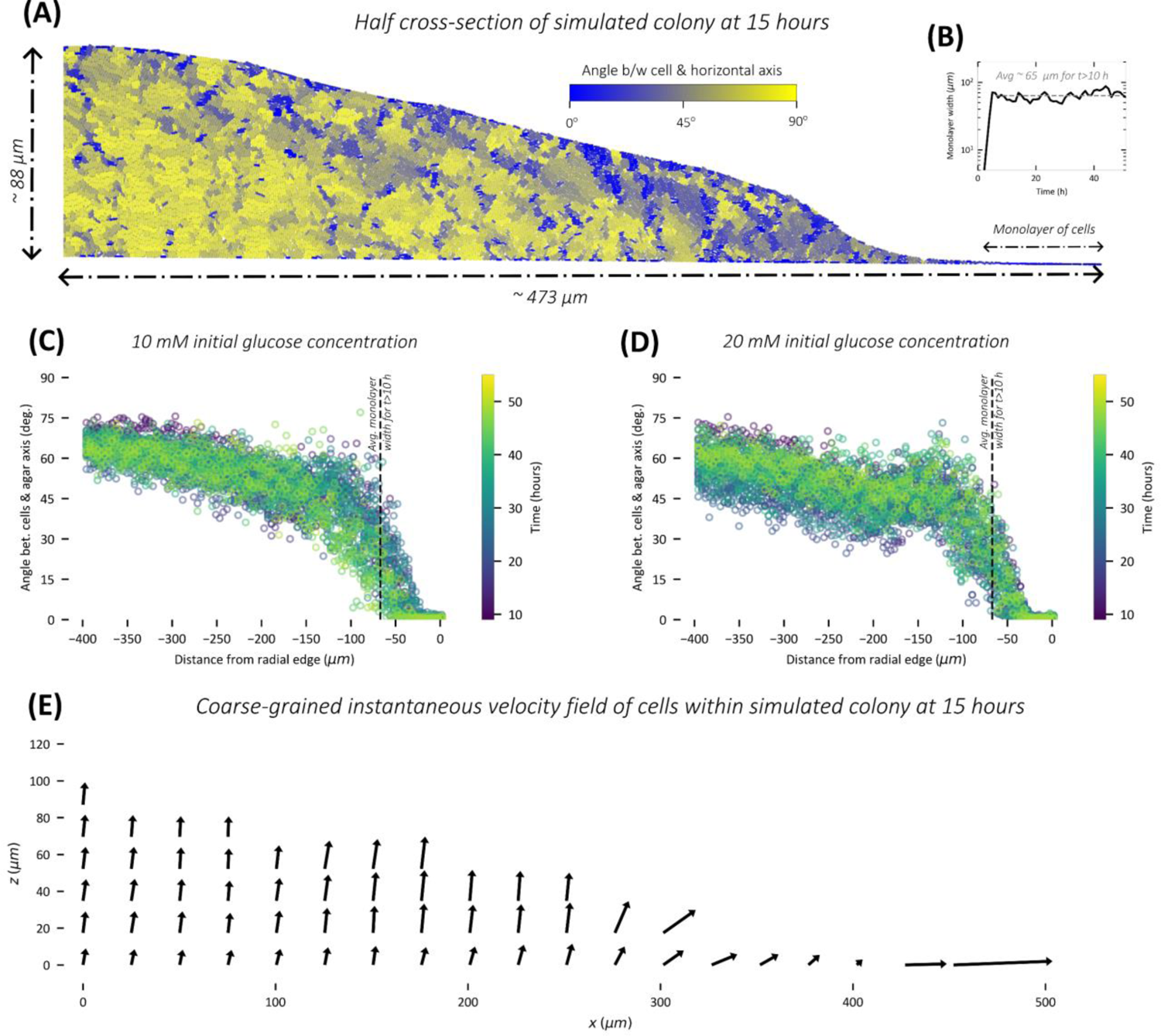
Orientation of cells within the colony. **(A)** Snapshot of cells within a 15 h old (1+1)-dimensional simulated colony with 10 mM initial glucose concentration. Cells are colored based on angle made with the horizontal axis (agar axis) where blue represents horizontal orientation and yellow represents vertical orientation. **(B)** Monolayer width (μm) of colony (averaged on both peripheral ends of colony) as a function of time for simulation with 20 mM initial glucose concentration. Angle made by cells with the horizontal axis (agar axis) plotted as a function of their distance from peripheral edge for **(C)** 10 mM initial glucose concentration and **(D)** 20 mM initial glucose concentration simulations. The angle plotted is averaged over the vertical sections of the colony at a particular distance from either peripheral edge of the colony. Zero degrees represent horizontal orientation, and 90 degrees represent vertical orientation. A distance of zero represents the peripheral edge and negative values indicates moving into the colony interior. **(E)** Snapshot of coarse-grained velocity field of cells within a 15 h old (1+1)-dimensional simulated colony with 10 mM initial glucose concentration.

**Extended Data Fig.5.**
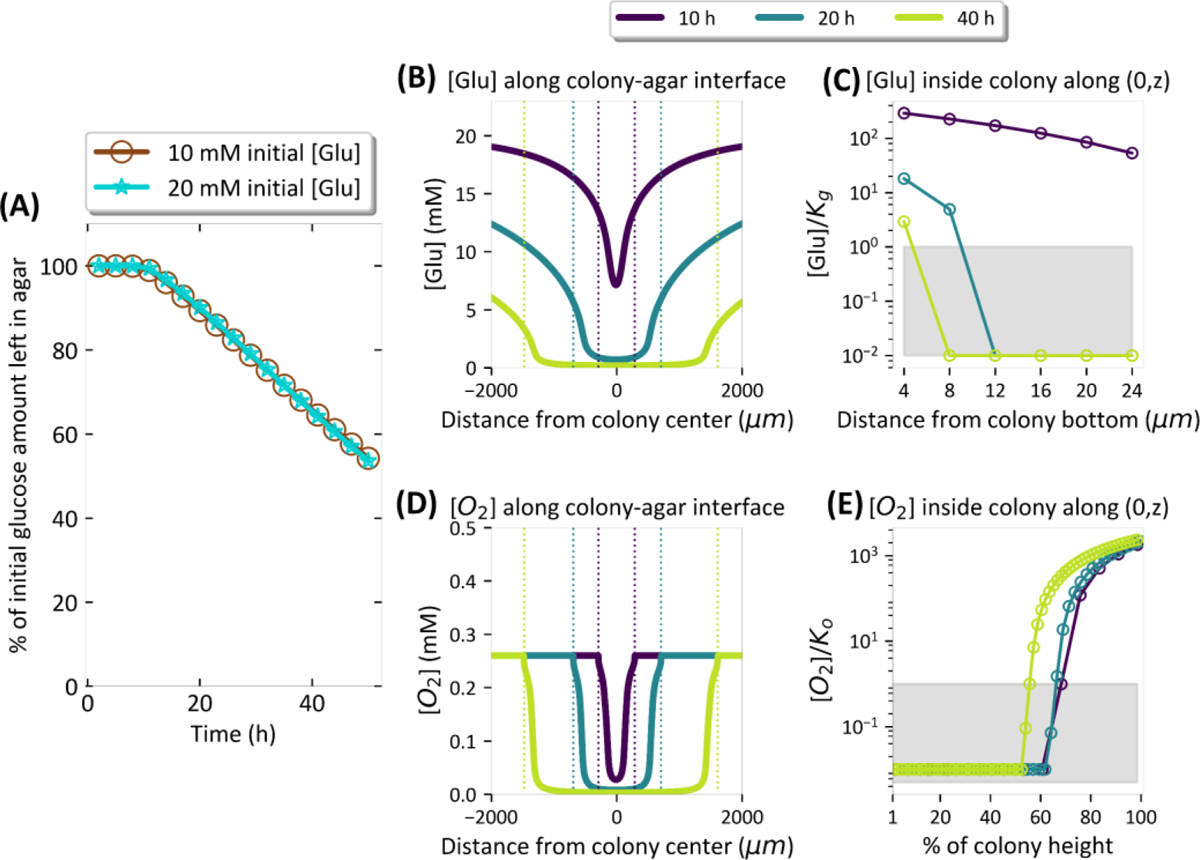
Spatiotemporal dynamics of glucose and oxygen along the agar surface and inside the colony. **(A)** The percentage of the initial amount of glucose (glucose concentration integrated over agar area) left in agar plotted as a function of time for simulations with 10 mM and 20 mM initial glucose concentration in agar. **(B)** The glucose concentration (mM) along the colony-agar interface plotted against the distance (μm) from colony center at 10 h, 20 h and 40 h stages of colony development (time coded by color). The colored vertical dotted lines indicate the colony edge at the corresponding stages of colony development. **(C)** Glucose concentration normalized by Kg along the central vertical axis of the colony plotted against the distance (μm) from colony bottom at 10 h, 20 h and 40 h stages of colony development (time coded by color). The grey shaded region represents concentrations below the Monod constant. **(D)** The oxygen concentration (mM) along the colony-agar interface plotted against the distance (μm) from colony center at 10 h, 20 h and 40 h stages of colony development (time coded by color). The colored vertical dotted lines indicate the colony edge at the corresponding stages of colony development. **(E)** The oxygen concentration normalized by Ko along the central vertical axis of the colony plotted against the distance from colony bottom expressed as percentage of colony height at 10 h, 20 h and 40 h stages of colony development (time coded by color). The grey shaded region represents concentrations below the Monod constant. Results shown here are for simulations with 20 mM initial glucose concentration.

**Extended Data Fig.6.**
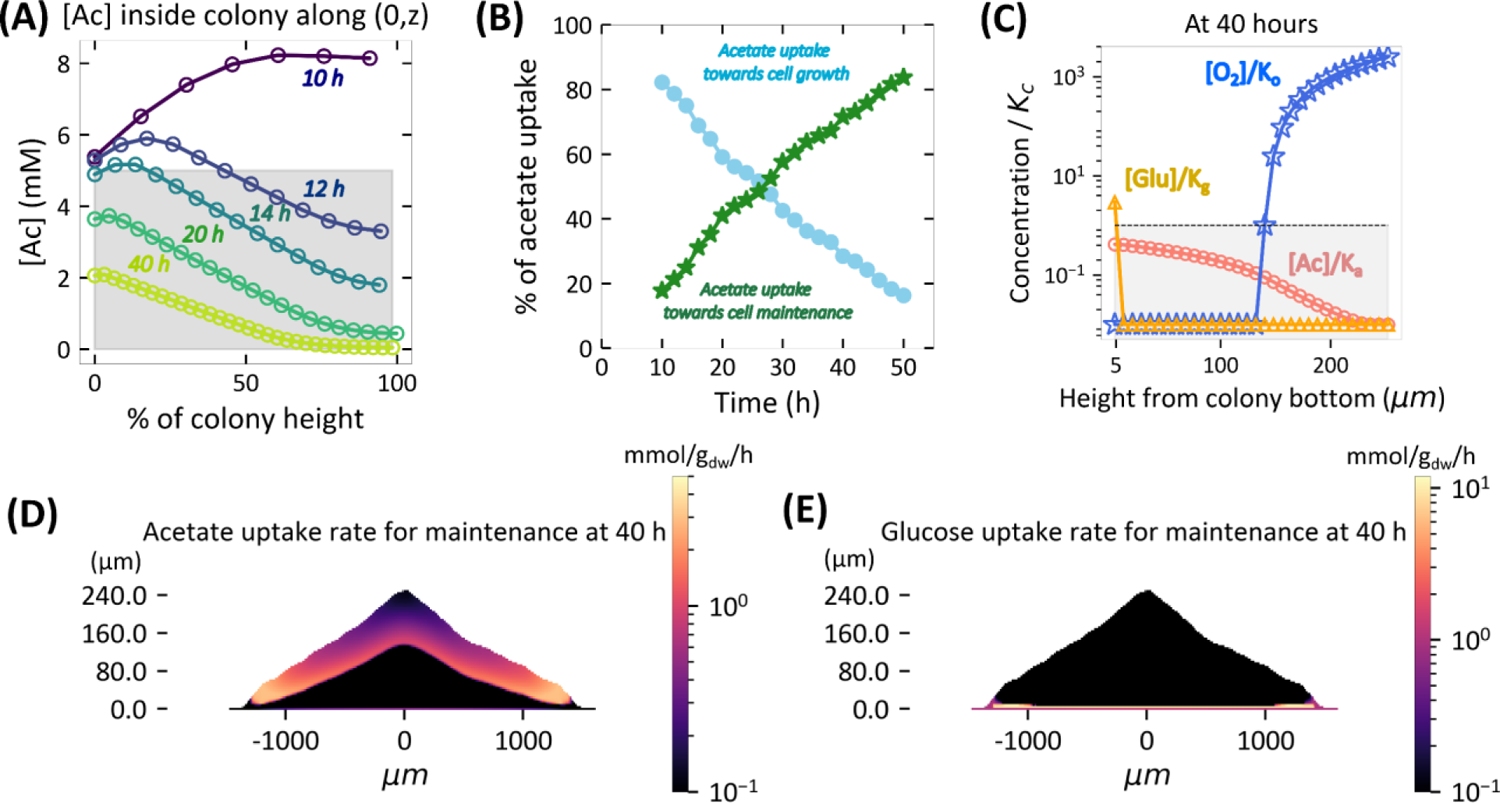
Spatiotemporal dynamics of acetate within colony and its role in cell maintenance. **(A)** Acetate concentration (mM) along the central vertical axis of the simulated colony plotted against the distance (μm) from colony bottom at 10 h, 12 h, 14 h 20 h and 40 h stages of colony development (time coded by color). The grey shaded region represents concentrations below the Monod constant. **(B)** The percentage of the acetate uptake flux (integrated over entire colony) towards cell growth (light blue) and towards maintenance (green) plotted against time (h) of colony development. **(C)** Glucose (yellow), oxygen (blue) and acetate (salmon) concentration normalized by respective Monod constants Kg, Ko, and Ka along the central vertical axis of the colony plotted against the vertical distance from colony bottom (μm) at 40 h stage of colony development. The grey shaded region indicates the concentration falling below the Monod constant. **(D)** Spatial distribution of acetate uptake rate for cell maintenance in mmol/gdw/h units (coded by color) within 40 h old colony is localized at the colony top near colony-air interface. **(E)** Spatial distribution of glucose uptake rate for cell maintenance in mmol/gdw/h units (coded by color) within 40 h old colony is localized at the colony bottom near colony-agar interface. Results shown here are for simulations with 20 mM initial glucose concentration.

**Extended Data Fig.7.**
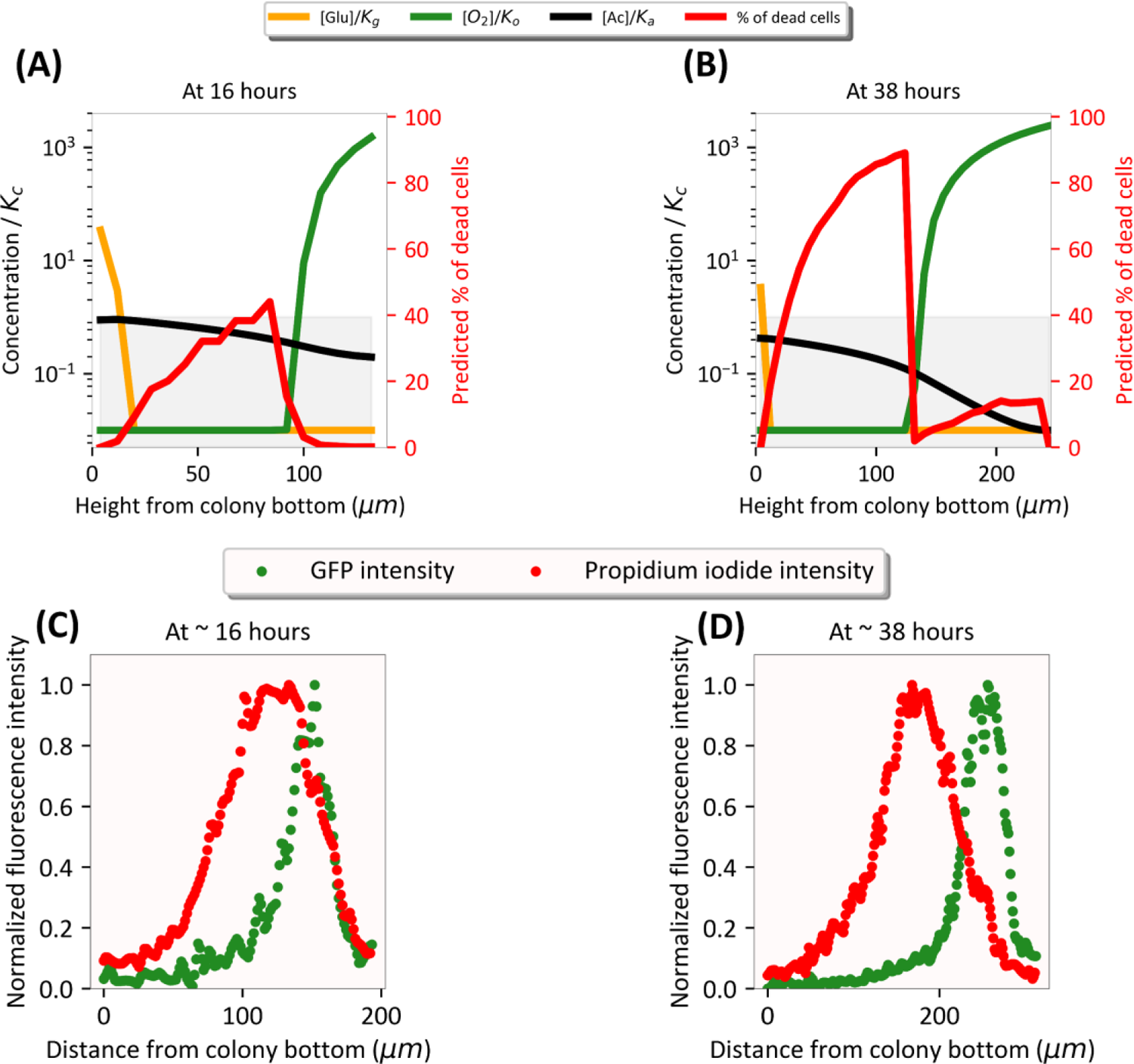
Death zone is localized to the colony center. **(A-B)** Left y-axis: Glucose (yellow), oxygen (green) and acetate (black) concentration normalized by respective Monod constants Kg, Ko, and Ka along the central vertical axis of the simulated colony plotted against the vertical distance from colony bottom (μm) at **(A)** 16 h stage and **(B)** 38 h of simulated colony development. Right y-axis: The predicted % of dead cells (red) along the central vertical axis of the simulated colony is plotted against the vertical distance from colony bottom (μm). **(C-D)** GFP intensity (in green) and propidium iodide fluorescence intensity (in red) normalized by the respective maximum intensities along the optical cross-section at the center of a **(C)** ∼ 16 h old and **(D)** ∼ 38 h old experiemental colony obtained using two-photon microscopy (See Methods). Simulation and experimental results presented here correspond to 20 mM initial glucose concentration.

**Extended Data Fig.8.**
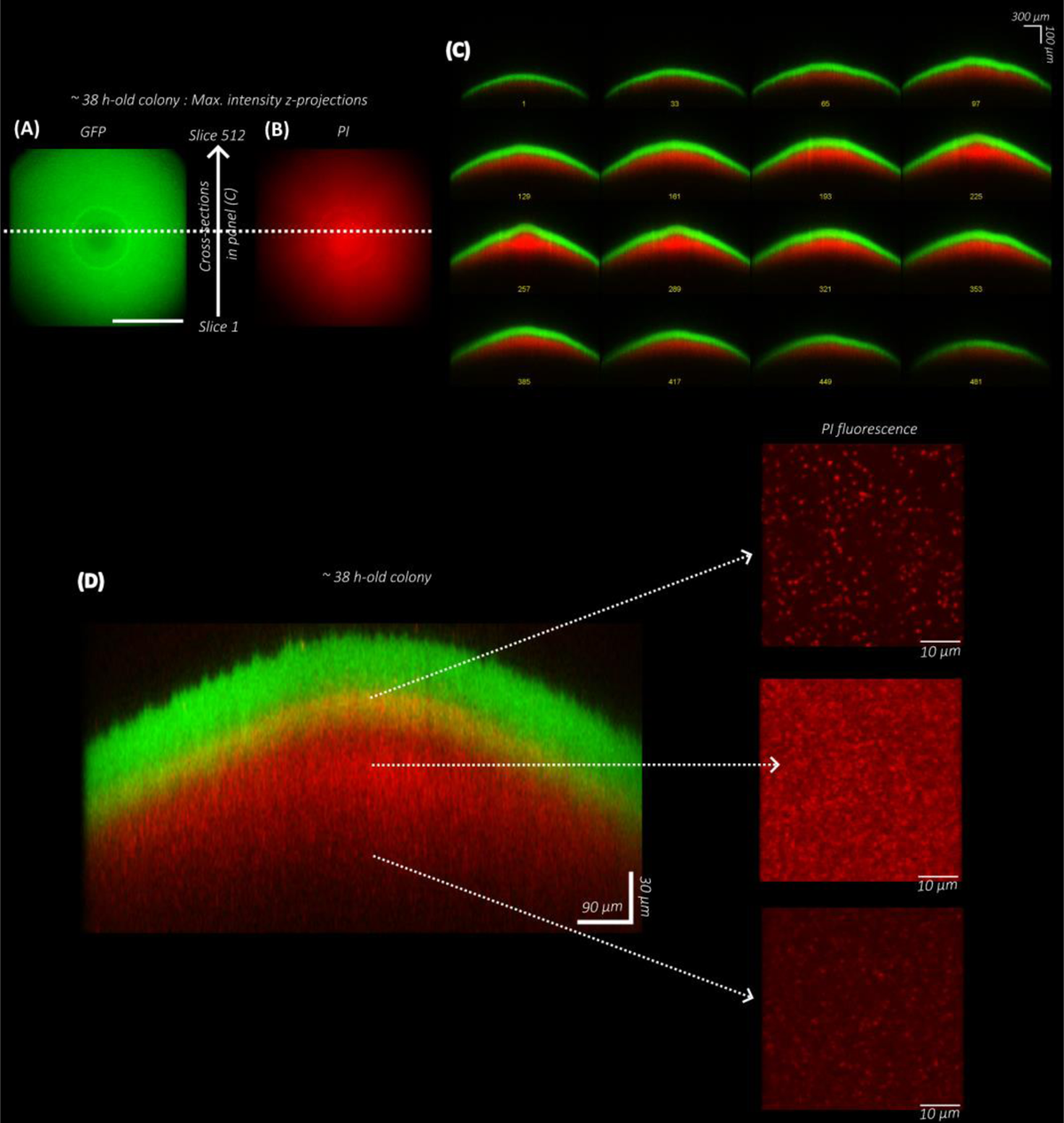
PI fluorescence along cross-sections of a 2 days-old colony. Maximum intensity z-projection of **(A)** GFP fluorescence intensity (in green) and **(B)** Propidium Iodide (PI) fluorescence (in red) of a ∼38 h old colony obtained using two-photon microscopy (see Methods). Scale bar in (A) represents 1000 μm. **(C)** Merged images of GFP fluorescence (in green) and Propidium iodide fluorescence (in red) along the various cross-sections of a ∼ 38 h-old colony. **(D)** Zoomed-in image of a central cross-section of a ∼38 h-old colony with a further magnified view of propidium iodide fluorescence at different heights. Colonies represented in panel **(C)** and **(D)** are from different experiments. For panel **(D)**, to get higher resolution images, a water immersion objective (25x/1 NA; Olympus, XLPLN25XSVMP2) was used. A thin layer of low-melt agarose was poured carefully at ∼ 37 °C onto the agar plate with the colony few minutes before imaging to permit the use of water immersion objective to image the colony from the top in an upright position.

**Extended Data Fig.9.**
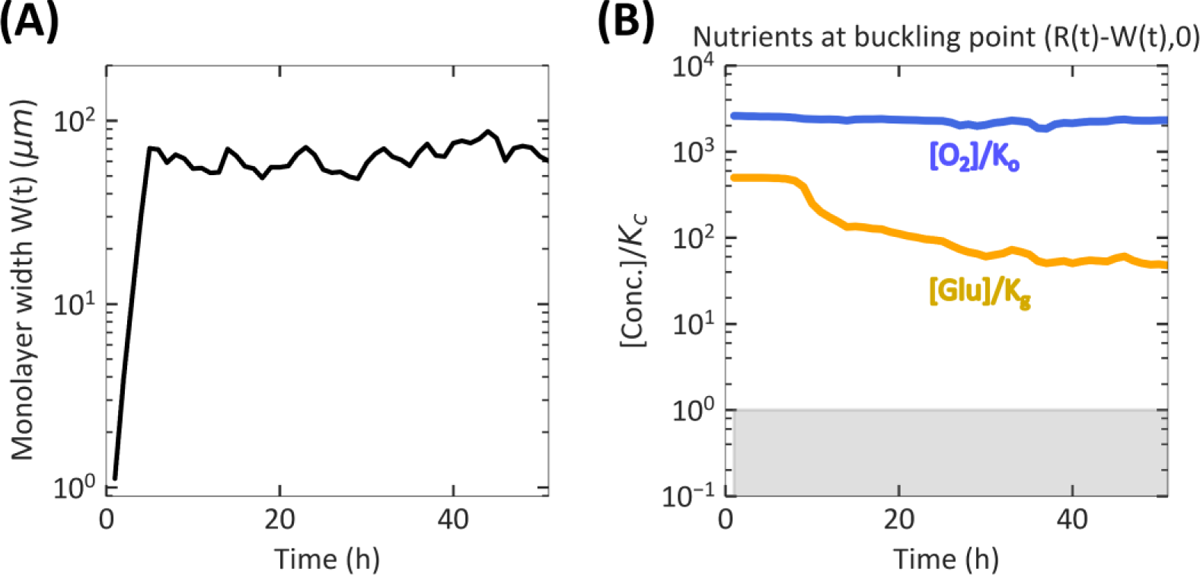
Nutrient concentrations are saturating at the buckling point of the colony thorughout the first 50 h. **(A)** The monolayer width (μm) of simulated colony plotted against time of colony development (h). **(B)** The concentration of glucose (yellow) and oxygen (blue) at the buckling point on the colony-agar interface i.e., the spot where the colony is no longer a monolayer is plotted against time of colony development (h). The grey shaded region represents nutrient concentrations below the corresponding Monod constant. Results shown here are for simulations with 10 mM initial glucose concentration.

**Extended Data Fig.10.**
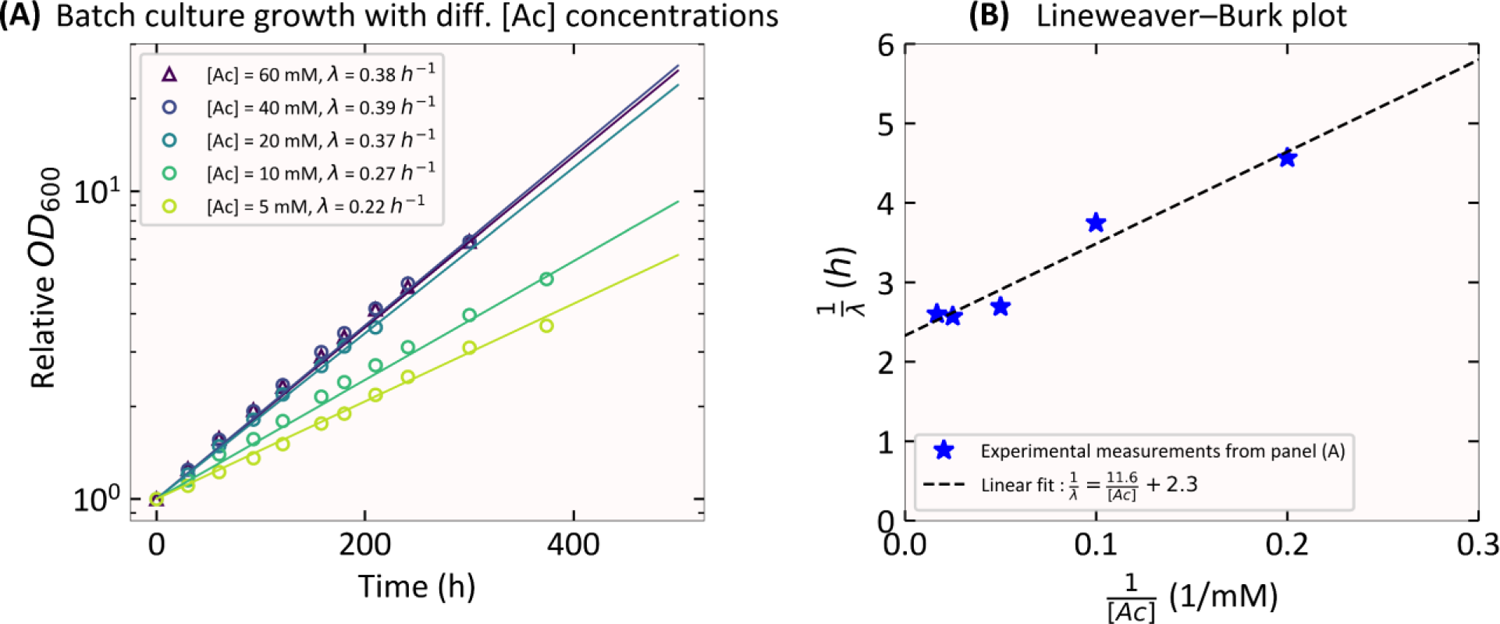
Growth of E. coli with acetate as the sole-carbon source. **(A)** Batch culture growth curves when EQ59 is grown with a defined concentration ranging from 5 mM to 60 mM of sodium acetate (coded by color) as the sole carbon source, 10 mM ammonium chloride and 112 mM phosphate buffered minimal medium. Batch culture growth is performed in a 37 °C water bath shaker (220 rpm). Cells from a glycerol stock stored at −80 °C are streaked on an LB agar plate and incubated overnight at 37 °C. A single colony from this plate is picked and inoculated into LB broth and grown in a shaking water bath for several hours at 37 °C as seed cultures. Seed cultures are then transferred into minimal medium with sodium acetate as the sole carbon source and grown overnight at 37 °C in a shaking water bath as pre-cultures. For the experimental cultures used to measure growth rate, overnight pre-cultures are diluted to around 0.01 to 0.02 OD600 in fresh minimal medium and grown at 37 °C in a shaking water bath as experimental cultures. **(B)** Lineweaver-Burk plot of reciprocal of growth rate vs. the reciprocal of acetate concentration (blue stars) where the growth rate for each condition is obtained by fitting data in panel **(A)** to an exponential function. Black dashed lines represent the best fit of data to a function of the form 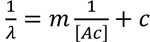. Assuming a Monod form for the growth rate dependence on acetate concentration i.e., 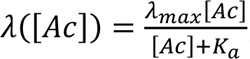, then note that 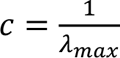 and 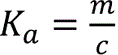. Based on the best linear fit (plotted in black dashed lines) *K_a_* ≈ 5 mM and λ_max_ ≈ 0.42 ℎ^−1^.

