## Supplementary Information for "Spatiotemporal development of growth and death zones in expanding bacterial colonies driven by emergent nutrient dynamics"

**4 Parameters** **9**

**5 Captions for Supplementary Videos** **11**

#### 1 An Agent-Based Model for Cell Activities

The agent-based component of our model to simulate the growth, division, and movement of individual bacterial cells within a colony growing on agar is adapted from Warren et.al. [10]. Here, we provide a brief overview of the key model components.

##### 1.1 Cell growth, division, and movement

We model an *E. coli* cell as a sphero-cylindrical agent, i.e., a cylinder with hemispherical caps on both ends. The length of the cylinder, $l$, elongates during cell growth, while the diameter, $w_0$, of the hemispherical caps (also referred to as the cell width) remains constant during the cell growth. We denote by $\vec{p}$ and $\vec{q}$ the position vectors of the centers of the two hemispheres, by $\ell = \|\vec{p} - \vec{q}\|$ the cell cylindrical length, and by $\vec{n} = (\vec{q} - \vec{p})/\ell$ the unit vector pointing from one center of hemisphere $\vec{p}$ to the other $\vec{q}$ and refer to it as the *director* of the cell. The center of mass for the cell is taken to be $\vec{r}_{\text{c}} = (\vec{p} + \vec{q})/2$.

The length $\ell = \ell(t)$ of a cell increases at the cell elongation rate $\tilde{\lambda}(t)$ with time $t$. The elongation rate is proportional to the mass growth rate $\lambda(t)$ of the cell, i.e., $\tilde{\lambda}(t) = \sigma\lambda(t)$, where the proportionality factor is $\sigma = \ln 3/\ln 2$ [10]. The mass growth rate of a cell at a particular time $t$ is calculated as $\lambda(t) = \lambda(\vec{r}_{\text{c}}(t), t)$, where $\vec{r}_{\text{c}}(t)$ is the center of the cell at time $t$ and $\lambda(\vec{r}, t)$ is the local growth rate at spatial point $\vec{r}$ and time $t$. The local growth rate is determined by the local concentrations of glucose, oxygen, and acetate (see Section 2.1 for details).

Each cell starts off with the same cylindrical length $\ell_0$ and grows with the elongation of its cylindrical length $\ell(t)$ governed by the growth equation,

$$\dot{\ell}(t) = \sigma\lambda(\vec{r}_{\text{c}}(t), t)\ell(t),$$

where $\dot{\ell}(t)$ denotes the time derivative of $\ell(t)$. Numerically, we update the cell length from time $t$ to $t + \Delta t$ by

$$\ell(t + \Delta t) = \ell(t) + \sigma\lambda(\vec{r}_{\mathsf{c}}(t), t)\ell(t)\Delta t.$$

Once the cylindrical length $\ell(t)$ increases by a $\Delta L$ amount, by the adder principle [7], the cell divides into two daughter cells. Upon division, the two daughter cells inherit the velocity and angular velocity from their mother cell with some fluctuations in their angular velocities.

The position and orientation of a cell changes according to its velocity $\vec{v}$ and angular velocity $\vec{\omega}$, which follow Newton's second law

$$M\frac{\partial \vec{v}}{\partial t} = \vec{F}^{\mathsf{net}} \qquad \text{and} \qquad I\frac{\partial \vec{\omega}}{\partial t} = \vec{T}^{\mathsf{net}}, \tag{1.1}$$

where $M$ and $I$ are the mass and moment of inertia of the cell, and $\vec{F}^{\mathsf{net}}$ and $\vec{T}^{\mathsf{net}}$ are the net force and net torque, respectively, exerted on that cell. A brief description of the forces exerted on a cell is provided in the next subsection. Once the net force and torque are calculated, following the numerical scheme used in [10], we use the velocity-Verlet algorithm [4] to update individual cell positions, orientations, velocities, and angular velocities based on Newton's law (Eq. [1.1]).

##### 1.2 Interaction forces

The forces and torques exerted on a cell within the simulated colony arise from the following four factors:

(1) **Cell-cell mechanical interaction:**
Following the implementation of Warren et.al. (2019) [10], two cells in contact with each other generate a contact force that is described by its components along the normal and tangential direction. The normal component of this force is the sum of the Hertz contact force component $\propto \sqrt{w_0}\delta_{\mathsf{cc}}^{3/2}$ (where $w_0$ is the cell-width, $\delta_{\mathsf{cc}}$ is the amount of overlap of the two cells), and the normal component of a dissipation force with magnitude $\propto \delta_{\mathsf{cc}}^{1/2} v_{\mathsf{cc},n}$ (where $v_{\mathsf{cc},n}$ is the normal component of the cell-cell relative velocity of the two cells). The tangential force component is taken to be the minimum of the tangential dissipation force component $\propto \delta_{\mathsf{cc}} v_{\mathsf{cc},t}$ (where $v_{\mathsf{cc},t}$ is the tangential component of the cell-cell relative velocity of the two cells ) and the Hertz contact force multiplied by the *dynamic* friction coefficient [10]. It should be noted that in high-density colonies such as those studied here, dissipation due to viscous drag is significantly less than the cellular friction force.

(2) **Cell-agar interaction** (if the cell is in contact with the agar surface):
The cell-agar interaction force is modeled as a contact force with elastic and frictional components similar to the cell-cell interaction force described above.

(3) **Cell-fluid interaction:**
The cell-fluid interaction force is modeled as a Stokes drag force which is proportional to the cell velocity.

(4) **Surface tension** (if the cell is at the colony-air boundary):
Surface tension between the air and liquid molecules coating a bacterial cell arises when the cell protrudes out on the top of bacterial colony, increasing the air-liquid interfacial area, thereby resulting in a restoring force exerted on the cell. The surface tension is implemented as a boundary force, i.e., as a force experienced by discrete cells at the colony boundary due to increased surface tension of the continuum liquid these cells are immersed in. As studied in Warren et al. (2019) [10], this individual cell-level implementation of surface tension can hold a large group of cells as a monolayer above the agar surface, until the pressure inside the expanding monolayer (due to friction against motion on agar surface) exceeds a critical level. At this critical level the pressure build up overcomes the surface tension resisting vertical protrusion of cells, thereby resulting in the 'buckling' of the monolayer of cells into multiple layers. This buckling of cells marks the transition from a 2D monolayer colony into a 3D colony.

#### 2 A Continuum Model for Metabolism

##### 2.1 Metabolic model to determine cell growth rate

The following three metabolic processes contribute to cell growth in our model:

(a) Aerobic growth on glucose: Cells consume glucose and oxygen, and produce biomass and excrete acetate.

$$\text{Glucose} + \text{Oxygen} \rightarrow \text{Biomass} + \text{Acetate};$$

(b) Anaerobic growth on glucose: Cells consume glucose in the absence of oxygen, and produce biomass and excrete acetate.

$$\text{Glucose} \rightarrow \text{Biomass} + \text{Acetate};$$

(c) Aerobic growth on acetate: Cells consume acetate and oxygen, and produce biomass.

$$\text{Acetate} + \text{Oxygen} \rightarrow \text{Biomass}.$$

We use appropriately modified Monod kinetic forms to determine the local cell growth rate as a function of the local concentrations of glucose $C_g = C_g(\vec{r}, t)$, acetate $C_a = C_a(\vec{r}, t)$, and oxygen $C_o = C_o(\vec{r}, t)$. First, we define for convenience,

$$\theta_g = \frac{C_g}{C_g + K_g}, \qquad \theta_a = \frac{C_a}{C_a + K_a}, \qquad \theta_o = \frac{C_o}{C_o + K_o}, \tag{2.1}$$

where $K_g$, $K_a$, and $K_o$ are the Monod constants corresponding to glucose, acetate and oxygen uptake by cells, respectively. We then determine the local cell growth rate $\lambda = \lambda(\vec{r}, t)$ at spatial point $\vec{r}$ in the colony and time $t$ by,

$$\lambda = \underbrace{\lambda_1(\lambda_{g,aer}, C_g)\theta_g\theta_o}_{\text{Aerobic growth on glucose}} + \underbrace{\lambda_2(\lambda_{g,ana}, C_g)(1-\theta_o)}_{\text{Anaerobic growth on glucose}} + \underbrace{\lambda_3(\lambda_{a,aer}, C_a)(1-\theta_g)\theta_o}_{\text{Aerobic growth on acetate}}. \tag{2.2}$$

Here, the factors $\theta_o$ and $(1 - \theta_o)$ are the weights for aerobic growth and anaerobic growth, respectively, and the factors $\theta_g$ and $1 - \theta_g$ are the weights for the growth on glucose and acetate, respectively. In a model which does not account for cell maintenance, the subscripted $\lambda$-terms would traditionally be defined using Monod kinetic forms, i.e.,

$$\lambda_1(\lambda_{g,aer}, C_g) = \lambda_{g,aer}\theta_g, \quad \lambda_2(\lambda_{g,ana}, C_g) = \lambda_{g,ana}\theta_g, \quad \lambda_3(\lambda_{a,aer}, C_a) = \lambda_{a,aer}\theta_a, \tag{2.3}$$

where, $\lambda_{g,aer}$, $\lambda_{g,ana}$, and $\lambda_{a,aer}$ are the maximum growth rates possible under aerobic growth on glucose, anaerobic growth on glucose, and aerobic growth on acetate, respectively. However, to incorporate the effects of cell maintenance we appropriately modify the local growth rates (see Section 2.3). Note that this growth model described by Eq. [2.2] serves as a tri-state switch, i.e., when both glucose and oxygen are abundant, aerobic growth on glucose will turn on; when oxygen is lacking, anaerobic growth on glucose will turn on; and when glucose is lacking, aerobic growth on acetate will turn on.

#### 2.2 Reaction-diffusion model for spatiotemporal dynamics of nutrients

The geometry of our model for a bacterial colony growing on hard agar is illustrated in Figure S1. The computational domain, $\Omega = (-L, L) \times (-L, L) \times (-a, b)$, includes the air region $\Omega_0$ (colored blue), the colony region $\Omega_1$ (colored salmon), and the agar region $\Omega_2$ (colored yellow). We denote by $\Gamma_{01}$ the interface separating the air region $\Omega_0$ and the colony region $\Omega_1$, by $\Gamma_{02}$ the interface separating the air region $\Omega_0$ and the agar region $\Omega_2$, and by $\Gamma_{12}$ the interface that separates the agar region $\Omega_1$ and colony region $\Omega_2$. The lateral and bottom faces of the agar region $\Omega_2$ are marked by $\Gamma_s$ and $\Gamma_b$, respectively. Since the bacterial colony grows with time $t$, all the air region $\Omega_0$, the colony region $\Omega_1$, the colony-air interface $\Gamma_{01}$, and the colony-agar interface $\Gamma_{02}$ depend on time $t$. Note that oxygen in the air region $\Omega_0$ diffuses into the bacterial colony and that glucose in agar diffuses into the colony and taken up by bacterial cells.

A system of reaction-diffusion partial differential equations (PDEs) for metabolite concentrations is used to model the spatiotemporal dynamics of metabolism within the colony. The concentrations of glucose $C_g = C_g(\vec{r}, t)$, acetate $C_a = C_a(\vec{r}, t)$, and oxygen $C_o = C_o(\vec{r}, t)$ are all defined spatially in both the colony region $\Omega_1$ which expands with time and the agar region $\Omega_2$ that is fixed throughout the simulation. For convenience, we shall also denote the colony and agar regions by $\Omega_+$ (same as $\Omega_1$) and $\Omega_-$ (same as $\Omega_2$), respectively.

**Glucose.** The reaction-diffusion equations for the glucose concentration are given by

$$\frac{\partial C_g}{\partial \mathrm{t}} = D_{g,-}\Delta C_g \qquad \text{in agar region } \Omega_-, \tag{2.4}$$

$$\frac{\partial C_g}{\partial \mathrm{t}} = D_{g,+}\Delta C_g - \rho Q_g \qquad \text{in colony region } \Omega_+. \tag{2.5}$$

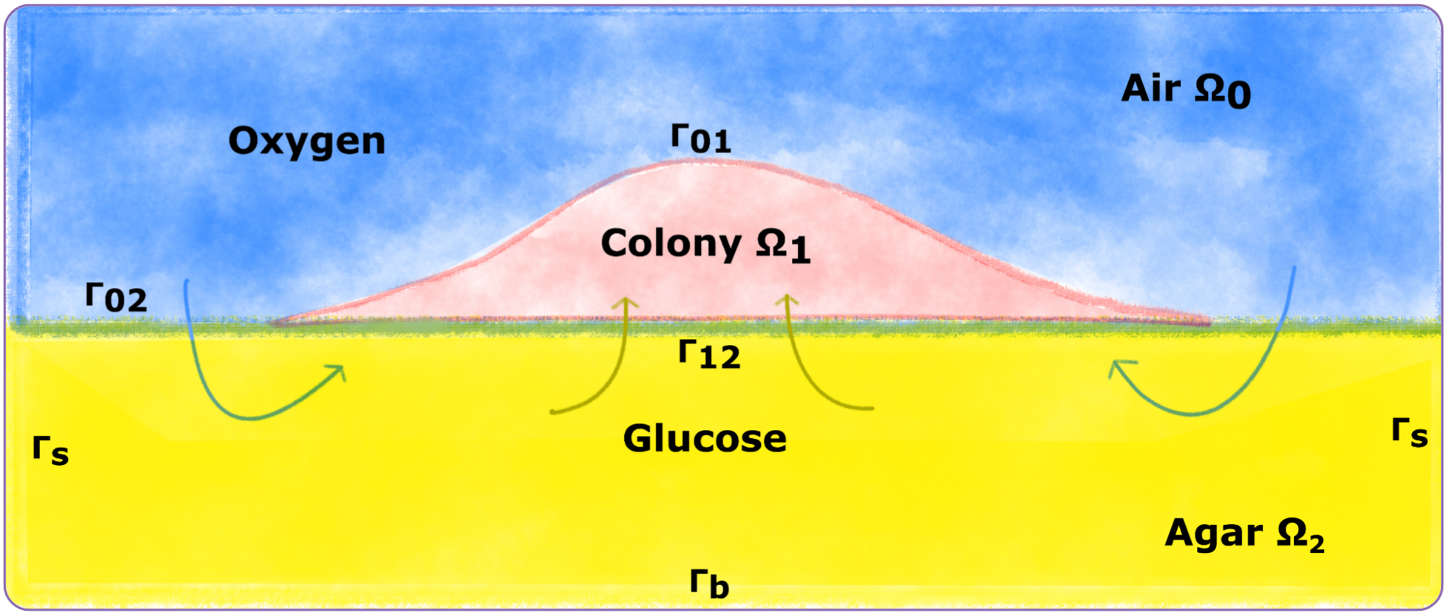

**SI Appendix Fig. S1:** An illustration of the computational domain and the different sub-regions involved in the simulations of colony growth. The overall region is $\Omega = (-L, L) \times (-L, L) \times (-a, b)$, where all $L$, $a$, and $b$ are positive numbers in the units of length. It is divided into the air region $\Omega_0$, colony region $\Omega_1$, and agar region $\Omega_2 = (-L, L) \times (-L, L) \times (-a, 0)$, respectively. The colony surface or colony-air interface $\Gamma_{01}$ separates the colony from air. The plane $z = 0$ in the system region is divided into two parts. One is the interface that separates the colony from agar, and is denoted by $\Gamma_{12}$. The other part, denoted $\Gamma_{02}$, separates the air from agar. Note that, since the bacterial colony grows with time $t$, the air region $\Omega_0$, the colony region $\Omega_1$, the colony-air interface $\Gamma_{01}$, and the colony-agar interface $\Gamma_{02}$, all depend on time $t$.

Here, $D_{g,-}$ and $D_{g,+}$ are the diffusion coefficients for glucose in agar and colony, respectively, $\rho$ is the local cell density within the colony, and $Q_g$ is the glucose consumption rate. We model such a rate with Monod kinetics:

$$Q_g = \underbrace{q_{g,aer}\lambda_1(\lambda_{g,aer}, C_g)\theta_g\theta_o}_{\text{Aerobic glucose consumption}} + \underbrace{q_{g,ana}\lambda_2(\lambda_{g,ana}, C_g)(1-\theta_o)}_{\text{Anaerobic glucose consumption}}, \qquad [2.6]$$

where $q_{g,aer}$ and $q_{g,ana}$ denote the specific glucose uptake fluxes under aerobic and anaerobic growth conditions, respectively, and the $\lambda_1$-term and $\lambda_2$-term are the local growth rates for aerobic growth on glucose and anaerobic growth on glucose, respectively.

Equations (2.4) and (2.5) are supplemented with the following interface conditions on the colony-agar interface $\Gamma_{12}$ that couple the glucose concentration $C_{g,-}$ from the agar region and $C_{g,+}$ from the agar region:

$$C_{g,-} = C_{g,+} \quad \text{and} \quad D_{g,+}\frac{\partial C_{g,+}}{\partial z} = D_{g,-}\frac{\partial C_{g,-}}{\partial z} \quad \text{on colony-agar interface } \Gamma_{12}.$$

The boundary conditions for the glucose concentration are

$$\frac{\partial C_g}{\partial n} = 0 \qquad \text{on } \Gamma_s \cup \Gamma_b \cup \Gamma_{02} \cup \Gamma_{01},$$

where $\partial/\partial n$ denotes the normal derivative.

Given an initial value of the glucose concentration, the system of these reaction-diffusion equations, together with the interface conditions and boundary conditions, determine uniquely the spatiotemporal dynamics of the glucose. In our simulations, we set the initial value of glucose to be a constant, $C_{g,0}$, in the agar region $\Omega_-$ but $0$ in the colony region $\Omega_+$.

**Oxygen.** The reaction-diffusion equations for the oxygen concentration are given by

$$\frac{\partial C_o}{\partial \text{t}} = D_{o,-}\Delta C_o \qquad \text{in agar region } \Omega_-, \qquad [2.7]$$

$$\frac{\partial C_o}{\partial \text{t}} = D_{o,+}\Delta C_o - \rho Q_o \qquad \text{in colony region } \Omega_+, \qquad [2.8]$$

where $D_{o,-}$ and $D_{o,+}$ are the diffusion coefficients for oxygen in agar and colony, respectively, and $Q_o$ is the oxygen consumption rate. We take the following form of this rate:

$$Q_o = \underbrace{q_{o,g}\lambda_1(\lambda_{g,aer}, C_g)\theta_g\theta_o}_{\text{Oxygen uptake during growth on glucose}} + \underbrace{q_{o,a}\lambda_3(\lambda_{a,aer}, C_a)(1-\theta_g)\theta_o}_{\text{Oxygen uptake during growth on acetate}}, \quad [2.9]$$

where $q_{o,g}$ and $q_{o,a}$ are the specific uptake fluxes of oxygen during the aerobic growth on glucose and that on acetate, respectively, and $\lambda_3(\lambda_{a,aer}, C_a)$ is the growth rate for aerobic growth on acetate, given in Eq. (2.3).

The interface conditions for the oxygen concentration on the colony-agar interface that couple the oxygen concentration $C_{o,-}$ in the agar and $C_{o,+}$ in the colony are given by

$$C_{o,-} = C_{o,+} \quad \text{and} \quad D_{o,+}\frac{\partial C_{o,+}}{\partial z} = D_{o,-}\frac{\partial C_{o,-}}{\partial z} \quad \text{on colony-agar interface } \Gamma_{12}.$$

The boundary conditions for the oxygen concentrations are,

$$C_o = C_{o,0} \qquad \text{on } \Gamma_{01} \cup \Gamma_{02},$$
$$\frac{\partial C_o}{\partial n} = 0 \qquad \text{on } \Gamma_s \cup \Gamma_b,$$

where $C_{o,0}$ is the boundary value of oxygen concentration which is taken to be a constant of 0.26 mM, approximately the concentration of oxygen in air. We set the initial oxygen concentration to be 0.26 mM in the agar region. Note that our choice for the boundary value and initial value of oxygen concentration assumes equilibration between air and agar.

**Acetate.** The reaction-diffusion equations for the acetate concentration are given by

$$\frac{\partial C_a}{\partial t} = D_{a,-}\Delta C_a \qquad \text{in agar region } \Omega_-, \quad [2.10]$$
$$\frac{\partial C_a}{\partial t} = D_{a,+}\Delta C_a + \rho P_a - \rho Q_a \qquad \text{in colony region } \Omega_+, \quad [2.11]$$

where $D_{a,-}$ and $D_{a,+}$ are the diffusion coefficients for acetate in agar and colony, respectively, $P_a$ is the acetate excretion rate during cell growth on glucose, and $Q_a$ is the acetate consumption rate during the cell growth on acetate. We model these rates using appropriate Monod kinetic forms as follows:

$$P_a = \underbrace{p_{a,aer}\lambda_1(\lambda_{g,aer}, C_g)\theta_g\theta_o}_{\text{Aceate production during aerobic growth}} + \underbrace{p_{a,ana}\lambda_2(\lambda_{g,ana}, C_g)(1-\theta_o)}_{\text{Acetate production during anaerobic growth}}; \quad [2.12]$$
$$Q_a = \underbrace{q_{a,aer}(1-\theta_g)\lambda_3(\lambda_{a,aer}, C_a)\theta_o}_{\text{Acetate consumption}}. \quad [2.13]$$

Here, $p_{g,aer}$ and $p_{g,ana}$ denote the specific excretion flux of acetate in the aerobic and anaerobic cell growth on glucose, respectively, $q_{a,aer}$ is the specific uptake flux of acetate during the cell growth on acetate, and all the subscripted $\lambda$-terms are growth rates for the three growth modes and are defined in Eq. (2.3).

The interface conditions for the acetate concentration on the colony-agar interface that couple the acetate concentration $C_{a,-}$ in the agar and $C_{a,+}$ in the colony are given by

$$C_{a,-} = C_{a,+} \quad \text{and} \quad D_{a,+}\frac{\partial C_{a,+}}{\partial z} = D_{a,-}\frac{\partial C_{a,-}}{\partial z} \quad \text{on colony-agar interface } \Gamma_{12}.$$

The boundary conditions for the acetate concentration are (Fig. S1)

$$\frac{\partial C_a}{\partial n} = 0 \qquad \text{on } \Gamma_{01} \cup \Gamma_{02} \cup \Gamma_s \cup \Gamma_b.$$

We set the initial acetate concentration to be $0$ in both the agar and colony regions.

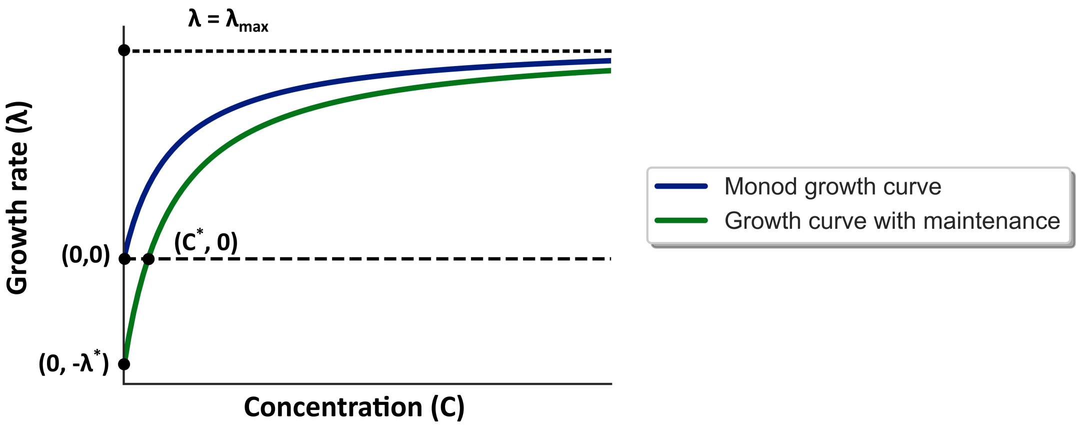

**SI Appendix Fig. S2:** Dependence of growth rate on nutrient concentration is modified to account for cell maintenance: cells grow only when the nutrient concentration is larger than a threshold value $C^*$.

#### 2.3 Model for cell maintenance

Even in the absence of growth, bacterial cells still need energy to perform maintenance activities. Such maintenance requires uptake of a carbon source. Three modes of cell maintenance, namely, maintenance on glucose under aerobic and anaerobic conditions and maintenance on acetate under aerobic conditions are included in our model. Further, with the inclusion of cell maintenance in our model, cells grow only when the nutrient concentration is larger than a threshold value $C^*$ (illustration in Fig. S2).

Let $q^0_{g,aer}$, $q^0_{g,ana}$, and $q^0_{a,aer}$ denote the maintenance rates for the three modes of maintenance, respectively. Further let,

$$\lambda^*_{g,aer} = \frac{q^0_{g,aer}}{q_{g,aer}}, \qquad \lambda^*_{g,ana} = \frac{q^0_{g,ana}}{q_{g,ana}}, \qquad \lambda^*_{a,aer} = \frac{q^0_{a,aer}}{q_{a,aer}}, \tag{2.14}$$

where $q_{g,aer}$, $q_{g,ana}$, and $q_{a,aer}$ are the specific uptake and excretion fluxes under various growth conditions which were introduced above in Eqns. (2.12) and (2.13). The total growth rate combined across the different modes has the same form as defined in Eq. (2.2). However, to additionally include the effect of maintenance, we modify the rates $\lambda_1$, $\lambda_2$, and $\lambda_3$ defined in Eq. (2.3) to be

$$\lambda_1(\lambda_{g,aer}, C_g) = \max\left\{(\lambda_{g,aer} + \lambda^*_{g,aer})\theta_g - \lambda^*_{g,aer},\, 0\right\}, \tag{2.15}$$

$$\lambda_2(\lambda_{g,ana}, C_g) = \max\left\{(\lambda_{g,ana} + \lambda^*_{g,ana})\theta_g - \lambda^*_{g,ana},\, 0\right\}, \tag{2.16}$$

$$\lambda_3(\lambda_{a,aer}, C_a) = \max\left\{(\lambda_{a,aer} + \lambda^*_{a,aer})\theta_a - \lambda^*_{a,aer},\, 0\right\}. \tag{2.17}$$

The threshold nutrient concentration (see Fig. S2) for the three growth modes, namely, $C^*_{g,aer}$, $C^*_{g,ana}$, and $C^*_{a,aer}$ are defined by

$$\lambda_1(\lambda_{g,aer}, C^*_{g,aer}) = 0, \qquad \lambda_2(\lambda_{g,ana}, C^*_{g,ana}) = 0, \qquad \lambda_3(\lambda_{a,aer}, C^*_{a,aer}) = 0.$$

This threshold concentration represents the local nutrient (glucose or acetate) concentration at which the maintenance rate corresponding to the particular mode can be sustained. Beyond these threshold concentrations, the cell exhibits growth under the corresponding growth mode. Solving for $C^*_{g,aer}$, $C^*_{g,ana}$, and $C^*_{a,aer}$, we obtain

$$C^*_{g,aer} = K_g \frac{\lambda^*_{g,aer}}{\lambda_{g,aer}}, \qquad C^*_{g,ana} = K_g \frac{\lambda^*_{g,ana}}{\lambda_{g,ana}}, \qquad C^*_{a,aer} = K_a \frac{\lambda^*_{a,aer}}{\lambda_{a,aer}}.$$

Consequently, we can rewrite [2.15]–[2.17] as

$$\lambda_1(\lambda_{g,aer}, C_g) = \begin{cases} (\lambda_{g,aer} + \lambda^*_{g,aer})\theta_g - \lambda^*_{g,aer}, & \text{if } C_g > C^*_{g,aer} \\ 0 & \text{if } C_g \leq C^*_{g,aer} \end{cases},$$

$$\lambda_2(\lambda_{g,ana}, C_g) = \begin{cases} (\lambda_{g,ana} + \lambda^*_{g,ana})\theta_g - \lambda^*_{g,ana}, & \text{if } C_g > C^*_{g,ana} \\ 0 & \text{if } C_g \leq C^*_{g,ana} \end{cases},$$

$$\lambda_3(\lambda_{a,aer}, C_a) = \begin{cases} (\lambda_{a,aer} + \lambda^*_{a,aer})\theta_a - \lambda^*_{a,aer}, & \text{if } C_a > C^*_{a,aer} \\ 0 & \text{if } C_a \leq C^*_{a,aer} \end{cases}.$$

The form of the overall local growth rate at a given spatial location is still the same as given in Eq. (2.2), i.e. ,

$$\lambda = \lambda_1(\lambda_{g,aer}, C_g)\theta_g\theta_o + \lambda_2(\lambda_{g,ana}, C_g)(1-\theta_o) + \lambda_3(\lambda_{g,aer}, C_a)(1-\theta_g)\theta_o. \quad [2.18]$$

Further, to account for the uptake of nutrients due to cell maintenance activities, we modify the uptake and excretion rates $Q_g$, $P_a$, $Q_a$, and $Q_o$, defined in Eqns. (2.6), (2.12), (2.13), and (2.9) to be

$$Q_g^M = Q_g + q^0_{g,aer}\tau(C_g, C^*_{g,aer})\,\theta_g\theta_o + q^0_{g,ana}\tau(C_g, C^*_{g,ana})\,(1-\theta_o), \quad [2.19]$$
$$P_a^M = P_a + p^0_{a,ana}\tau(C_g, C^*_{g,ana})\,(1-\theta_o), \quad [2.20]$$
$$Q_a^M = Q_a + q^0_{a,aer}\tau(C_a, C^*_{a,aer})\,(1-\theta_g)\theta_o, \quad [2.21]$$
$$Q_o^M = Q_o + q^0_{o,g}\tau(C_g, C^*_{g,aer})\,\theta_g\theta_o + q^0_{o,a}\tau(C_a, C^*_{a,aer})\,(1-\theta_g)\theta_o, \quad [2.22]$$

where, $q^0_{o,g}$ and $q^0_{o,a}$ denote the uptake rate of oxygen during the maintenance on glucose and acetate respectively, and

$$\tau(x,a) = \begin{cases} x/a & \text{if } x \le a, \\ 1 & \text{if } x > a. \end{cases}$$

The above $M$-superscripted rates replace $Q_g$, $P_a$, $Q_a$, and $Q_o$ in Eqns. (2.5), (2.8), and (2.11) to incorporate metabolite uptake and acetate excretion during cell-maintenance.

#### 2.4 Model for tracking nutrient starvation and predicting cell death within colony

The instantaneous uptake flux by a cell for maintenance relative to the maintenance rate corresponding to each of the three maintenance modes, $M^{cell}_{g,aer}(t)$, $M^{cell}_{g,ana}(t)$ and $M^{cell}_{a,aer}(t)$, are defined as

$$M^{cell}_{g,aer}(t) = \tau(C_g, C^*_{g,aer})\theta_g\theta_o, \quad [2.23]$$
$$M^{cell}_{g,ana}(t) = \tau(C_g, C^*_{g,ana})(1-\theta_o), \quad [2.24]$$
$$M^{cell}_{a,aer}(t) = \tau(C_a, C^*_{a,aer})(1-\theta_g)\theta_o. \quad [2.25]$$

Note that $q^0_{g,aer}\tau(C_g, C^*_{g,aer})\theta_g\theta_o$, $q^0_{g,ana}\tau(C_g, C^*_{g,ana})(1-\theta_o)$ and $q^0_{a,aer}\tau(C_a, C^*_{a,aer})(1-\theta_g)\theta_o$ are the absolute maintenance uptake rates corresponding to the respective mode of maintenance (as appearing in [2.19] and [2.21]). A relative maintenance uptake rate value being $x$ (where $0 \le x \le 1$) for a particular maintenance mode indicates that the nutrient uptake rate of the cell meets $100\,x$% of the corresponding maintenance rate.

We introduce a quantity termed the *deficit* of a cell, $D^{cell}(t)$, to denote the magnitude of a cell's inability to meet the carbon maintenance rate, which is defined by

$$D^{cell}(t) = \max\left(1 - \left(M^{cell}_{g,aer}(t) + M^{cell}_{g,ana}(t) + M^{cell}_{a,aer}(t)\right),\ 0\right), \quad [2.26]$$

where $M^{cell}_{g,aer}(t) + M^{cell}_{g,ana}(t) + M^{cell}_{a,aer}(t)$ represents the relative carbon uptake flux towards maintenance summed across all three maintenance modes. Thus $D^{cell}(t)$ describes the instantaneous starvation state of a cell where a value of $1$ represents complete starvation whereas a value of $0$ represents no starvation, and a value between 0 and 1 represents that the maintenance flux is only met partially. Then, the carbon starvation duration, denoted by $\gamma^{cell}(T)$, for a particular cell at time $T$ of colony development is defined by

$$\gamma^{cell}(T) = \int_{\text{Birth time}}^{T} D^{cell}(t)\,dt. \quad [2.27]$$

To predict the probability of cell death based on starvation duration it is necessary to differentiate between aerobic starvation duration and anaerobic starvation duration. This is due to our experimental measurements (Fig. 7B of main text) which show that death rate under anaerobic carbon starvation $(2.32\ d^{-1})$ is roughly 10-fold higher than the death rate under aerobic carbon starvation $(0.21\,d^{-1})$. Thus, the instantaneous aerobic deficit $D_{aer}^{cell}(t)$ and anaerobic deficit $D_{ana}^{cell}(t)$ at time $t$ for a particular cell are defined as

$$D_{aer}^{cell}(t) = \begin{cases} \text{max}\left(1 - \left(M_{g,aer}^{cell}(t) + M_{a,aer}^{cell}(t)\right),\ 0\right) & \text{if } M_{g,aer}^{cell}(t) > \epsilon \text{ or } M_{a,aer}^{cell}(t) > \epsilon \text{ (i.e., in aerobic condition),} \\ 0 & \text{else,} \end{cases} \quad [2.28]$$

$$D_{ana}^{cell}(t) = \begin{cases} \text{max}\left(1 - M_{g,ana}^{cell}(t),\ 0\right) & \text{if } M_{g,aer}^{cell}(t) < \epsilon \text{ and } M_{a,aer}^{cell}(t) < \epsilon \text{ (i.e., in anaerobic condition),} \\ 0 & \text{else.} \end{cases} \quad [2.29]$$

Here, $\epsilon$ is a number which acts as a threshold to determine whether the cell is under aerobic or anaerobic starvation, i.e., if the relative maintenance uptake rate through either aerobic maintenance mode is less than $\epsilon$, then the cell is classified to be under anaerobic condition and vice versa. In our simulations $\epsilon$ is chosen to be 0.01.

Given the instantaneous aerobic and anaerobic deficit of a cell, the aerobic carbon starvation duration $\gamma_{aer}^{cell}(T)$ and anaerobic carbon starvation duration $\gamma_{ana}^{cell}(T)$ at time $T$ of colony development are defined by

$$\gamma_{aer}^{cell}(T) = \int_{\text{Birth time}}^{T} D_{aer}^{cell}(t)\,dt, \quad [2.30]$$

$$\gamma_{ana}^{cell}(T) = \int_{\text{Birth time}}^{T} D_{ana}^{cell}(t)\,dt. \quad [2.31]$$

Once the aerobic carbon duration $\gamma_{aer}^{cell}(T)$ and anaerobic carbon starvation $\gamma_{ana}^{cell}(T)$ of a particular cell at time $T$ is known, the probability of the cell being dead at time $T$ of colony development is determined by the following equation:

$$\text{Death probability of cell at time } T = \text{max}\left(1 - e^{-0.21\gamma_{aer}^{cell}(T)},\ 1 - e^{-2.32\gamma_{ana}^{cell}(T)}\right) \quad [2.32]$$

where $0.21\ d^{-1}$ is the death rate under aerobic carbon starvation and $2.32\ d^{-1}$ is the death rate under anaerobic carbon starvation as motivated by our batch culture glucose starvation experiments (Fig. 7B of main text).

### 3 Numerical Methods and Computer Implementation

#### 3.1 A (1+1)-dimensional approximation

In order to computationally simulate colony expansion lasting two days by which the radial dimension is in the order of millimeters and the vertical dimension is hundreds of micrometers, we approximate our underlying colony system with a (1+1)-dimensional geometry. The cells in the colony occupy a two-dimensional region with one dimension being along the colony-agar interface and the other dimension being the one perpendicular to the agar surface. Further, the components of forces experienced by cells along the excluded dimension are set to zero. This two-dimensional setting allows study of both radial and vertical colony expansion in a computationally tractable manner.

#### 3.2 Overall time iteration

We use our hybrid discrete-continuum model to simulate a colony growing on hard agar from time $t = 0$ to a final simulation time $t = T_{final}$ in hours. Such simulations are done through a time iteration with a uniform macro time step $\Delta t$ and a total of $N_{overall}$ time steps, where $N_{overall}\Delta t = T_{final}$. Initially, we distribute glucose uniformly in the agar region with a constant glucose concentration $C_{g,0}$. To begin the simulation, we place one cell at the center of the agar surface. The initial velocity and angular velocity of this cell are set to be zero. In each time iteration, we simulate the colony growth for a time period $\Delta t$. Each such individual iteration with the time step $\Delta t$ consists of the following main steps:

(1) Generate the macroscopic boundary of colony $\Gamma_{01}$ (Fig. S1) by thresholding the cell density.
(2) Update the concentrations of glucose, oxygen, and acetate $C_g$, $C_o$, and $C_a$, respectively, by solving the system of reaction-diffusion equations for these concentrations using $N_{conc}$ iterations with a small time step $dt$, where, $N_{conc}dt = \Delta t$.
(3) Update the local cell growth rate.

(4) Use the agent-based model to simulate the cell growth and division, compute the cell interaction forces and torques, and simulate the cell movement using the velocity-Verlet method [4], for all the cells in the colony with a small time step $\delta t$ and a total number of iterations $N_{cell}$ with $N_{cell}\delta t = \Delta t$.

We refer readers to Warren et al. (2019) [10] for further details on model aspects involving the agent-based treatment of growth, division, and movement of individual cells within the colony.

##### 3.3 Numerical solution to reaction-diffusion equations

Given the colony boundary $\Gamma_{01}$ (Fig. S1) and concentrations $C_g$, $C_o$, and $C_a$ of glucose, oxygen, and acetate, respectively, in both the agar region $\Omega_-$ and colony region $\Omega_+$ at time $t = n\Delta t$ for some $n$ $(0 \le n < N_{overall})$, we solve the system of reaction-diffusion equations for the time period from $t = n\Delta t$ to $t = (n+1)\Delta t$ using $N_{conc}$ iterations with each iteration having a time step $dt$ to obtain all the concentrations at time $t = (n+1)\Delta t$. The initial values of the concentrations are those at the time $t = n\Delta t$ (i.e., the concentrations from the previous macro time step $n\Delta t$).

We use different methods of numerical discretization for solving the partial differential equations involved in the agar and colony regions. We use the Crank–Nicholson scheme for the time-dependent diffusion equation in the agar region $\Omega_-$ with the time step size $dt$, and use the forward Euler method to discretize the reaction-diffusion equation in the colony region $\Omega_+$ with a smaller time step size $dt/N_{colony}$ for some large integer $N_{colony}$. The two parts are coupled through the interface conditions, i.e., the continuity of the concentration and its flux across the agar-colony interface. We adopt the nested finite difference grids as in Warren et al. (2019) [10], with a fine grid covering the colony region and a region in agar beneath the colony, and several levels of coarser grids covering the rest of the agar region. For regular grid points inside the colony region (those grid points that are away from the colony surface), we use the standard five-point finite-difference scheme to discretize the diffusion operator. For those irregular points, the grid points inside the colony but are close to the colony surface, we introduce ghost grid points outside but close to the colony, and use interpolation to assign the values of concentration at these grid points, and then discretize the diffusion operator at irregular grid points inside the colony. In the agar region, we discretize the diffusion operator using the standard five-point finite-difference scheme at the grid points that are not on the interface between the fine and coarse grids, and using the standard scheme and interpolation for the interface grid points.

For each time step, we use the multigrid method to solve the system of algebraic linear equations resulting from the time and spatial discretization in the agar region, and use the forward Euler method to find steady-state solutions to the reaction-diffusion equations in the colony region. Since we take the solution at the previous step as our initial condition, the number of forward Euler steps required for convergence is small.

**Algorithm 1** Algorithm for numerically solving reaction-diffusion equation

`Input` : Model and numerical parameters (see Section 4), colony shape determined by the boundary $\Gamma_{01}$ (Fig. S1), concentrations $C_g$, $C_o$ ,and $C_a$ from the previous macro time step.
`Output` : Concentrations $C_g$, $C_o$, and $C_a$ in the agar and colony regions.
`Initialization:` Set $n = 0$.
**while** $n < N_{conc}$ **do**
  • Use the multigrid method to solve the linear system of algebraic equations resulting from the time and spatial discretization of the reaction-diffusion equations in the agar region $\Omega_-$.
  • Update the concentrations $C_g$, $C_o$, and $C_a$ at the colony-agar interface using the interface conditions.
  • Use the forward Euler method to solve the reaction-diffusion equations for the steady state concentrations $C_g$, $C_o$, and $C_a$ in the colony region $\Omega_+$, with the time step $dt/N_{colony}$;
  • Update $n \leftarrow n + 1$.
**end while**

#### 4 Parameters

The parameters involved in the metabolic model are listed below in Table 1. Numerical parameters involved in solving the partial differential reaction-diffusion equations for nutrient concentrations are listed separately in Table 2. The model and numerical parameters for the agent-based discrete simulations for cellular mechanical interaction forces are similar to values used in Warren et al. (2019) [10].

**Table 1:** Parameters for metabolic and reaction-diffusion model.

| Symbol | Description | Value | Units | Source (or) Rationale for choice |
|---|---|---|---|---|
| $\lambda_{g,aer}$ | Maximum batch culture growth rate aerobically on glucose | 0.9 | $h^{-1}$ | This study |
| $\lambda_{g,ana}$ | Maximum batch culture growth rate anaerobically on glucose | 0.6 | $h^{-1}$ | This study |
| $\lambda_{a,aer}$ | Maximum batch culture growth rate aerobically on acetate | 0.4 | $h^{-1}$ | This study |
| $q_{g,aer}$ | Specific flux for glucose during aerobic growth on glucose | 11 | $mmol/g_{DW}$ | [9] |
| $q_{g,ana}$ | Specific flux for glucose during anaerobic growth on glucose | 28 | $mmol/g_{DW}$ | [9] |
| $q_{a,aer}$ | Specific flux for acetate during aerobic growth on acetate | 33 | $mmol/g_{DW}$ | Equiv. to $q_{g,aer}$ on a per C-atom basis |
| $q_{o,g}$ | Specific flux for oxygen during aerobic growth on glucose | 22 | $mmol/g_{DW}$ | [1] |
| $q_{o,a}$ | Specific flux for oxygen during aerobic growth on acetate | 22 | $mmol/g_{DW}$ | [1] |
| $p_{a,aer}$ | Specific flux of acetate excretion during aerobic growth on glucose | 3 | $mmol/g_{DW}$ | [2] |
| $p_{a,ana}$ | Specific flux of acetate excretion during anaerobic growth on glucose | 16 | $mmol/g_{DW}$ | [8] |
| $q^0_{g,aer}$ | Maintenance rate for glucose in aerobic condition | 1 | $mmol/g_{DW}/h$ | [9] |
| $q^0_{g,ana}$ | Maintenance rate for glucose in anaerobic condition | 10 | $mmol/g_{DW}/h$ | [9] |
| $q^0_{a,aer}$ | Maintenance rate for acetate in aerobic condition | 3 | $mmol/g_{DW}/h$ | Equiv. to $q^0_{g,aer}$ on a per C-atom basis |
| $q^0_{o,g}$ | Uptake rate for oxygen during maintenance on glucose in aerobic condition | 2 | $mmol/g_{DW}/h$ | Approximated as $q_{o,g} \cdot \frac{q^0_{g,aer}}{q_{g,aer}}$ |
| $q^0_{o,a}$ | Uptake rate for oxygen during maintenance on acetate in aerobic condition | 2 | $mmol/g_{DW}/h$ | Approximated as $q_{o,a} \cdot \frac{q^0_{a,aer}}{q_{a,aer}}$ |
| $p^0_{a,ana}$ | Acetate excretion rate during anaerobic maintenance on glucose | 10 | $mmol/g_{DW}/h$ | $\approx q^0_{g,ana}$ (Ref. [6] 1 Glu $\rightarrow$ 1 Acetate + 1 Ethanol + 2 Formate) |
| $K_g$ | Monod constant for glucose | 20 | $\mu M$ | [5] |
| $K_a$ | Monod constant for acetate | 5 | $mM$ | This study (see Extended Data Fig. 10) |
| $K_o$ | Monod constant for oxygen | 0.1 | $\mu M$ | [3] |
| $C_{o,0}$ | Boundary value of oxygen concentration | 260 | $\mu M$ | [3] |
| $D_{g,+}$ | Diffusion coefficient for glucose in agar | 740 | $\mu m^2/s$ | [3] |
| $D_{g,-}$ | Diffusion coefficient for glucose in colony | 110 | $\mu m^2/s$ | [10] : $D_{g,+} \approx \phi D_{g,-}$ with $\phi \approx 0.15$ |
| $D_{a,+}$ | Diffusion coefficient for acetate in agar | 1100 | $\mu m^2/s$ | [3] |
| $D_{a,-}$ | Diffusion coefficient for acetate in colony | 165 | $\mu m^2/s$ | [10] : $D_{a,+} \approx \phi D_{a,-}$ with $\phi \approx 0.15$ |
| $D_{o,+}$ | Diffusion coefficient for oxygen in agar | 2500 | $\mu m^2/s$ | [3] |
| $D_{o,-}$ | Diffusion coefficient for oxygen in colony | 375 | $\mu m^2/s$ | [10] : $D_{o,+} \approx \phi D_{o,-}$ with $\phi \approx 0.15$ |

**Table 2:** Numerical parameters and cell-centric parameters.

| Symbol | Description | Value | unit |
|---|---|---|---|
| $L$ | Half-Length of agar region (Fig. S1) | 12160 | $\mu m$ |
| $a$ | Depth of agar region (Fig. S1) | 8192 | $\mu m$ |
| $b$ | Height of air region (Fig. S1) | 400 | $\mu m$ |
| $N_x$ | Number of grid points in the $x$ direction | 1520 | No unit |
| $N_{z\text{c}}$ | Number of grid points in the $z$ direction in colony | 100 | No unit |
| $N_{z\text{a}}$ | Number of grid points in the $z$ direction in agar | 512 | No unit |
| $h_{\text{grid,c}}$ | Spatial grid size in colony region | 4 | $\mu m$ |
| $M_{agar}$ | Number of grid levels in agar region (Multi-nested grid approach [10]) | 3 | No unit |
| $\Delta t$ | Macro-time step for simulations (see Section 3.2) | 0.01 | h |
| $dt$ | Time step while solving reaction-diffusion equations (see Section 3.2) | 0.00004 | h |
| $\delta t$ | Time step to simulate cell growth and movement due to forces (see Section 3.2) | 0.00001 | h |
| $N_{colony}$ | Maximum number of iterations for forward-Euler method to solve PDEs in colony region (see Section 3.3) | 10000 | No unit |
| $w_0$ | Diameter of hemispherical caps of a cell | 1 | $\mu m$ |
| $\Delta L$ | Length added to a cell before division occurs | 3 | $\mu m$ |
| $\rho_{cell}$ | Cell dry weight per cell volume of a cell | 0.25 | $pg/\mu m^3$ |

#### 5 Captions for Supplementary Videos

**Supplementary Video 1**. 3D rendering from confocal microscopy images of a $\sim$ 20 h old EQ59 colony grown on a 1.5 %(w/v) minimal media agar plate prepared with 20 mM glucose, 10 mM ammonium chloride, and 112 mM phosphate buffered saline.

**Supplementary Video 2**. 3D rendering from confocal microscopy images of a $\sim$ 53 h old EQ59 colony grown on a 1.5 %(w/v) minimal media agar plate prepared with 20 mM glucose, 10 mM ammonium chloride, and 112 mM phosphate buffered saline.

**Supplementary Video 3**. Spatiotemporal dynamics of glucose concentration (in units of $K_g$) during simulated colony development in the colony region and the rectangular sub-portion of agar region immediately beneath colony.

**Supplementary Video 4**. Spatiotemporal dynamics of oxygen concentration (in units of $K_o$) during simulated colony development in the colony region and the rectangular sub-portion of agar region immediately beneath colony.

**Supplementary Video 5**. Spatiotemporal dynamics of acetate concentration (in units of $K_a$) during simulated colony development in the colony region and the rectangular sub-portion of agar region immediately beneath colony.

**Supplementary Video 6**. Spatiotemporal dynamics of cell growth rate ($h^{-1}$) during simulated colony development.

**Supplementary Video 7**. Spatiotemporal dynamics of cell growth rate ($h^{-1}$) from (Top) aerobic glucose metabolism, (Middle) anaerobic glucose metabolism, (Bottom) aerobic acetate metabolism during simulated colony development.

**Supplementary Video 8**. Spatiotemporal dynamics of maintenance flux ($mmol/g_{dw}/h$) from (Top) glucose metabolism, (Bottom) acetate metabolism during simulated colony development.

**Supplementary Video 9**. Spatiotemporal dynamics of carbon starvation within colony during simulated colony development.

**Supplementary Video 10**. Spatiotemporal dynamics of the death zone within colony during simulated colony development.
